# *MSH2* knock-down shows CTG repeat stability and concomitant upstream demethylation at the *DMPK* locus in myotonic dystrophy type 1 human embryonic stem cells

**DOI:** 10.1101/2020.09.25.313197

**Authors:** Silvie Franck, Lise Barbé, Simon Ardui, Yannick De Vlaeminck, Joke Allemeersch, Dominika Dziedzicka, Claudia Spits, Fien Vanroye, Pierre Hilven, Geoffrey Duqué, Joris R. Vermeesch, Alexander Gheldof, Karen Sermon

## Abstract

Myotonic dystrophy type 1 (DM1) is caused by expansion of a CTG repeat in the *DMPK* gene, where expansion size and somatic mosaicism correlates with disease severity and age of onset. While it is known that the mismatch repair protein MSH2 contributes to the unstable nature of the repeat, its role on other disease-related features, such as CpG methylation upstream of the repeat, is unknown. In this study, we investigated the effect of an *MSH2* knock-down (MSH2KD) on both CTG repeat dynamics and CpG methylation pattern in human embryonic stem cells (hESC) carrying the DM1 mutation. Repeat size in MSH2 wild type (MSH2WT) and *MSH2KD* DM1 hESC was determined by PacBio sequencing and CpG methylation by bisulfite massive parallel sequencing. We found stabilization of the CTG repeat concurrent with a gradual loss of methylation upstream of the repeat in MSH2KD cells, while the repeat continued to expand and upstream methylation remained unchanged in MSH2WT control lines. Repeat instability was re-established and biased towards expansions upon *MSH2* transgenic re-expression in MSH2KD lines while upstream methylation was not consistently re-established. We hypothesize that the hypermethylation at the mutant DM1 locus is promoted by the MMR machinery and sustained by a constant DNA repair response, establishing a potential mechanistic link between CTG repeat instability and upstream CpG methylation. Our work represents a first step towards understanding how epigenetic alterations and repair pathways connect and contribute to the DM1 pathology.

## Introduction

Myotonic dystrophy type 1 (DM1 [MIM: 160900]) is caused by an unstable CTG repeat expansion in the 3’ UTR of the *DMPK* gene and mainly affects the neuronal and muscular systems^1^. Patients have an expanded and unstable CTG repeat tract sizing between 50 to 6500 repeats whereas unaffected individuals have a short and stable tract ranging from 5 to 37 CTG repeats^2,3^. Unusual DNA structures in the repetitive tract are suggested to serve as mutagenic intermediates explaining why repeat instability occurs whenever single stranded DNA is formed both in a replication-dependent or independent manner^4–6^. A mosaic pattern of different CTG repeat lengths is observed across different tissues, in which the most affected, including skeletal muscle, heart and brain, show the largest somatic expansions^7^. This somatic instability is also observed in human cell cultures^8–11^. Ongoing CTG expansions in somatic tissues promotes disease severity and progression, while intergenerational instability leads to the phenomenon known as genetic anticipation^11–14^. DM1 offspring usually exhibit longer repeat lengths than the affected transmitting parent and show symptoms earlier in life^2,15^. Anticipation in DM1 displays a clear parent-of-origin effect, as the repeat length tends to be longer in offspring from affected mothers, which explains the almost exclusive occurrence of the most severe congenital form of DM1, CDM1, with maternal transmission^16^. In the *DMPK* gene, the CTG repeat is embedded in a CpG-rich region with two CCCTC-binding factor sites (CTCF1 and CTCF2) located up- and downstream of the repeat^17,18^. In a human primary fibroblast model, Cho *et al*. (2005) demonstrated that the CTCF protein is unable to bind its target DM1 sequence when it is CpG methylated^17^. We recently demonstrated that methylation of the CpG island upstream of the CTG repeat, containing a CTCF binding site (CTCF1), is strongly associated with CDM1^19^.

The link between the mismatch repair pathway (MMR) components MSH2 and MSH3 (forming the MutSβ complex) and expansion-biased trinucleotide repeat (TNR) instability is well established^5,20^. DM1 mice deficient for Msh2^21,22^ and DM1 human induced pluripotent stem cells (hiPSCs) with knocked-down *MSH2*^23^ were reported to display reduced repeat expansions or even repeat contractions. An Msh3 depletion resulted in halting somatic repeat instability in transgenic mice^24^. Furthermore, MSH6 also forms a complex with MSH2, MutSα, but this complex seems to have only a minor role in TNR instability^5,24^. Msh6 depletion even caused a higher somatic repeat instability in transgenic mice, explained by the fact that the higher availability of MSH2 to form complexes with MSH3 led to higher instability ^24^. We previously demonstrated a stabilization of the CTG repeat in human embryonic stem cells (hESCs) carrying the DM1 mutation upon their differentiation into a non-affected cell type, osteogenic progenitor-like-cells (OPLs), concomitant with a marked natural decrease of MMR components expression, in particular MSH2^9^.

A number of reports have shown inconsistent findings regarding a correlation of methylation with CTG instability, showing either an absence^25^, a significant tendency towards a correlation^19,26,27^ or even a strong correlation^28^ between the CTCF1 methylation status and repeat length or instability. Other authors even hypothesized that the expanded repeat was the causative agent which induced upstream methylation^29^. The link between active CTG instability and upstream methylation at the DM1 allele remains unclear.

Using *MSH2* knock-down (MSH2KD) DM1 hESC we studied the relationship between upstream CpG methylation, CTG repeat instability and the mismatch repair machinery at the DM1 allele. We observe a statistically significant decrease in repeat expansions as well as a trend towards repeat contractions after *MSH2* knock-down whereas the CTG repeat in MSH2WT clonal lines continued to expand. Concurrently, the upstream CpG region (CTCF1) lost DNA methylation upon *MSH2* knock-down, which might suggest a molecular link between TNR instability and CpG methylation upstream of the CTG repeat. Re-establishing *MSH2* expression in MSH2KD clonal lines induced repeat expansions but not a systematical CpG methylation upstream of the CTG repeat, suggesting that MSH2 is linked to the maintenance but not the *de novo* methylation of this locus.

## Results

We generated three single-cell (SC) clonal *MSH2* knock-down lines (VUB03KD-SC1, VUB03KD-SC2, VUB03KD-SC3) from hESC line VUB03-DM1 and one (VUB19KD-SC1) from VUB19-DM1, using CRISPR/Cas9 genome editing (**Figure 1A**). The guide RNA of the CRISPR/Cas9 system targets exon 3 of the *MSH2* gene, which encodes the DNA binding domain of the protein. The mutations introduced by the CRISPR/Cas9 system were assessed through Sanger sequencing and massive parallel sequencing of a 371 nt PCR product spanning the gRNA target sequence (**Figure S1**). The introduced mutations can be found in **Figure S2A**. VUB03KD-SC2 and VUB03KD-SC3 show an apparently homozygous in-frame mutation where possibly the second allele is a large deletion and is probably not covered by the PCR (**Figure S2A, B**). VUB03KD-SC1 has a stop gain and frameshift mutation for both alleles and VUB19KD-SC1 has two different frameshift mutations for both alleles (**Figure S2A, B**). Two non-DM1 lines were genome-edited as controls resulting in two *MSH2* knock-down lines from VUB02 (VUB02KD-SC1, VUB02KD-SC2) and two from VUB06 (VUB06KD-SC1, VUB06KD-SC2) (**Figure 1A**). The introduced mutations can be found in **Figure S3A, B**. Each MSH2KD line is the result of a clonal expansion of a single hESC after genome editing. This ensured a homogeneous MSH2KD phenotype in all cells of a clonal line as well as a homogeneous starting point regarding CTG repeat size and CpG methylation, in contrast to the mosaic CTG repeat size and CpG methylation in the original hESC line. We also prepared single cell clones from the starting MSH2WT lines, generating 5 clones: VUB03WT-SC1, VUB03WT-SC2 and VUB03WT-SC3 for cell line VUB03-DM1 and VUB19WT-SC1 and VUB19WT-SC2 for VUB19-DM1 (**Figure 1A**). Clonal lines were also derived from non-DM1 control lines: two MSH2WT clonal lines (VUB02WT-SC1, VUB02WT-SC2) from VUB02 and one (VUB06WT-SC1) from VUB06, (**Figure 1A**).

**Figure 1.**
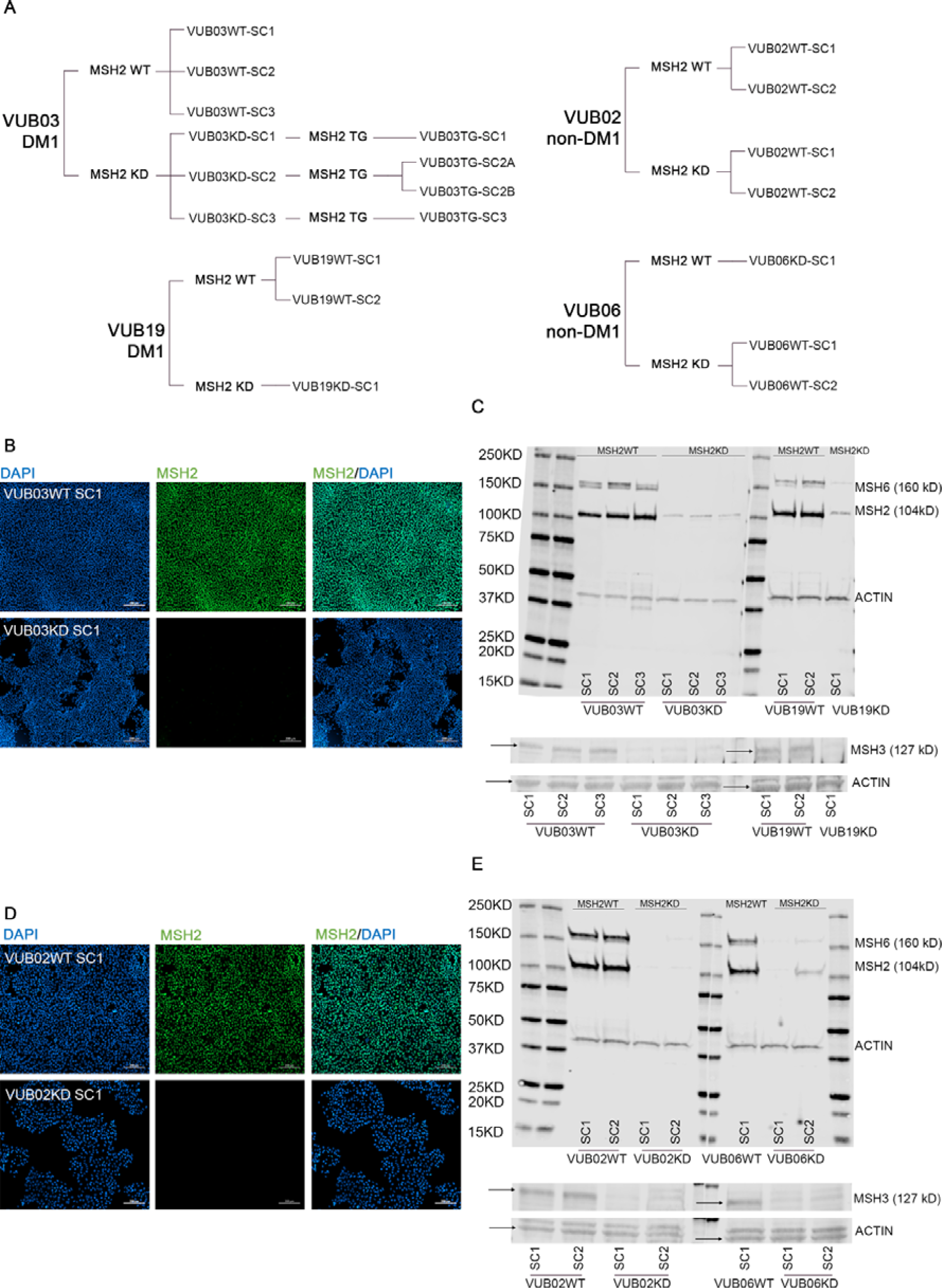
Characterization of the *MSH2KD* DM1 human embryonic stem cell lines. (A) The embryonic stem cell collection and their derived MSH2WT, MSH2KD and MSH2TG clonal cell lines. VUB03-DM1 and VUB19-DM1 were derived from embryos carrying DM1. VUB02 and VUB06 are non-affected cell lines with a CTG repeat size of 5 and 12 repeats. (B) Immunocytochemistry for MSH2 (GFP) showing a positive signal in VUB03WT-SC1 and the absence of MSH2 in VUB03KD-SC1. Immunocytochemistry images and quantification of WB results of all clonal lines are presented in **Figure S2, S4**. (C) Western blot showing MSH2 and MSH6 protein expression in DM1 lines, and a detail of the WB for MSH3. The complete MSH3 WB is shown in **Figure S4A**. ACTIN was used as endogenous protein loading control. (D) Immunocytochemistry for MSH2 (GFP) showing a positive signal in VUB02WT-SC1 and the absence of MSH2 in VUB02KD-SC1. Immunocytochemistry images and quantification of WB results of all clonal lines are presented in **Figure S3, S4**. (E) Western blot of control lines. The complete WB for MSH3 can be found in **Figure S4B**. Abbreviations: GFP: Green Fluorescence, SC: single cell clonal line, KD: MSH2 knock-down, WT: MSH2 wild type

All MSH2KD lines showed a significantly reduced MSH2 protein expression, as demonstrated by immunocytochemistry **(Figure 1B and D, S2C, S3C)** and Western blot **(Figure 1C and E, S4)**. MSH2 protein is reduced 6 to 10-fold in our VUB03KD and VUB19KD lines when compared to their respective MSH2WT controls VUB03WT-SC1 and VUB19WT-SC2 as quantified by WB band intensity (quantification in **Figure S4C**). It has been previously reported that MSH2 is an obligatory component of the MMR system, and its inactivation leads to post-translational down-regulation of its heterodimerizing partners MSH3 and MSH6 ^31^. MSH6 and MSH3 protein levels also significantly decreased in MSH2KD cell lines, as expected, due to loss of stability when unable to complex with MSH2 **(Figure S4E-H)**^30^.

We determined the CTG repeat size at *DMPK* in all DM1 MSH2KD and MSH2WT lines by small-pool PCR followed by PacBio sequencing^32^, at passages 4, 12 and 20 after cloning of the lines. All MSH2WT lines show CTG repeat instability, with a progressive increase of repeat size with time in culture (**Figure 2, right panel)** (Jonckheere-Terpstra Test *p*<0.05, **Table S2**). Over 16 passages, the median size of the expanded allele in the three VUB03-DM1 MSH2WT lines (median CTG size: 732, 684 and 710 repeats, respectively, at passage 4) increased with 215 repeats in VUB03WT-SC1 and VUB03WT-SC3 and 185 repeats in VUB03WT-SC2. The lines of VUB19-DM1 started with a smaller repeat size (median CTG size: 375 repeats), but also increased with 66 repeats in VUB19WT-SC1 and 84 repeats in VUB19WT-SC2 over the consecutive passages. In contrast, significantly less repeat size variability is seen in all MSH2KD lines **(Figure 2, right panel)**, together with the tendency to slowly contract (Jonckheere-Terpstra Test *p*<0.05) (**Table S2**). A decrease in median size of the CTG expansion with 34, 23 and 24 repeat units in lines VUB03KD-SC1, VUB03KD-SC2 and VUB03KD-SC3 respectively and a decrease of 15 units in VUB19KD-SC1 was observed. This change from repeat expansion towards contractions upon *MSH2* knock-down is consistent with other published findings^21,23,33^. A detailed overview of the CTG repeat size heterogeneity in all lines is shown in **Table S3**.

**Figure 2.**
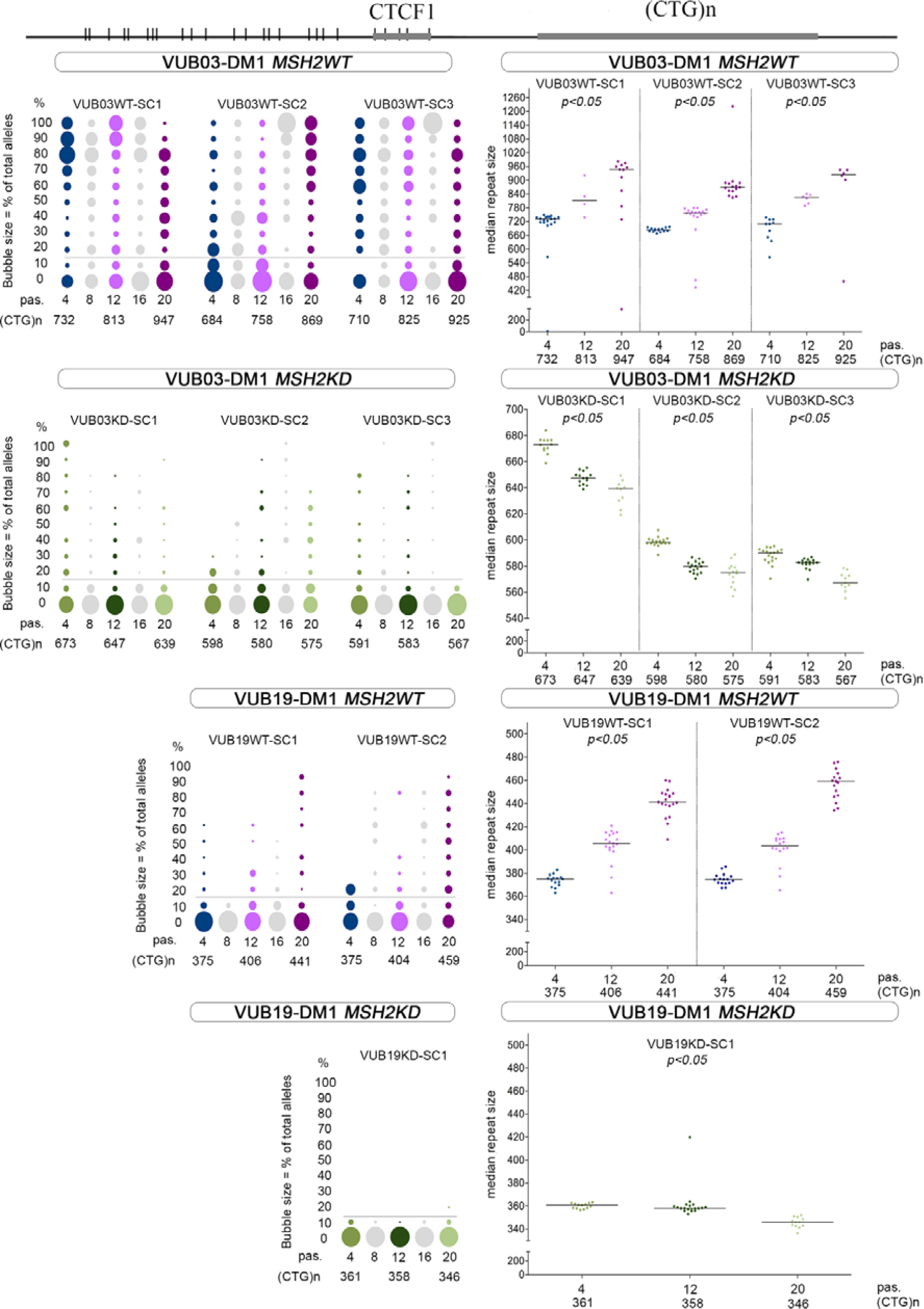
*DMPK* CTG repeat contraction and loss of methylation in the upstream CpG rich region in *MSH2KD* clonal lines. The top of the graph represents the *DMPK* CTG repeat region, with the 25 upstream CpG sites (vertical bars) and the CTCF1 site. The methylation levels are shown on the left and CTG repeat length distributions are shown on the right per clonal line of VUB03-DM1 and VUB19-DM1 from passage 4 to passage 20. The X-axis shows the passage number and median CTG repeat size. Methylation is shown for 25 CpG sites and a total of 100 epi-alleles were analyzed per cell passage. Bubble size is relative to the number of epi-alleles with the indicated percentage of CpG methylation, as previously described ^19^. The horizontal grey line represents the threshold for non-methylated alleles and is established on methylation levels in non-DM1 individuals as described in Barbé et al. (2017) ^19^. All alleles above the threshold are considered as methylated alleles. In the right panel, each dot represents the median (most frequently present) CTG repeat size of one low input LongAmp PCR reaction, spanning the repeat. The horizontal line represents the median for that clonal cell line and passage number. Abbreviations: pas.: passage

Subsequently, we determined the CpG methylation at the flanking sites of the CTG repeat at the *DMPK* locus in all MSH2KD DM1 and isogenic control lines, at multiple time points between passages 4 to 20. We used bisulfite treated massive parallel sequencing of the flanking regions to study 25 CpG sites upstream of the repeat and 11 located downstream. The upstream region includes 5 CpG’s located in CTCF1 (**Figure 2, left panel)** ^19^. At passage 4, all MSH2WT clonal lines of VUB03-DM1 had methylated epi-alleles (VUB03WT-SC1: 84%, VUB03WT-SC2: 48% and VUB03W-SC3: 83%, **Table S4**) and retained this methylation profile over culture for 16 passages (p=0.646, **Table S5**). In contrast, all VUB03-DM1 MSH2KD lines showed a rapid and nearly complete, statistically significant loss of upstream methylation (p=0.025, **Table S5**). By passage 4 after *MSH2* knock-down, only 39%, 12% and 19% of the epi-alleles remained methylated in VUB03KD-SC1, VUB03KD-SC2 and VUB03KD-SC3 respectively (**Table S4**) and these lines retained their low methylated state over culture (p=0.448, **Table S5**).

For VUB19-DM1, MSH2WT clonal lines were largely unmethylated at passage 4 (9% for VUB19WT-SC1 and 25% for VUB19WT-SC2, **Table S4**), but progressively gained methylation over time in culture (p=0.021, **Table S5**) concurrent with an increase in CTG repeat size. At early passages, there was no significant difference in methylation between VUB19-DM1 MSH2WT and MSH2KD lines. However, at passage 20 VUB19-DM1 MSH2WT lines were significantly more methylated than the MSH2KD line at the same passage (p=0.005, **Table S5**), indicating an increase of methylation in the MSH2WT sublines while MSH2KD lines retained low methylation (0%, **Table S4**) over culture (p=1, **Table S5**). The methylation pattern at the CpG region downstream of the repeat, spanning the CTCF2 binding site, remained unchanged in all lines (MSH2WT and MSH2KD) and irrespective of the time in culture (**Figure S5**).

The methylation status of non-DM1 MSH2WT (VUB02WT-SC1, VUB02WT-SC2, VUB06WT-SC1) and MSH2KD (VUB02KD-SC1, VUB02KD-SC2, VUB06KD-SC1, VUB06KD-SC2) lines was determined by Sanger sequencing following bisulfite treatment. The results showed, as expected, no methylation at any of the CpG sites both upstream (**Figure S6**) and downstream (**Figure S7**) of the repeat.

As a remarkable stabilization of the repeat together with demethylation of the upstream region was observed upon *MSH2* knock-down, we next re-established *MSH2* expression in knocked-down clonal lines. *MSH2* was re-expressed using lentiviral transduction at passage 12 of the three VUB03-DM1 MSH2KD lines. The lines underwent a new cloning event, resulting in 4 *MSH2* transgenic (TG) clonal lines (VUB03TG-SC1 from VUB03KD-SC1, VUB03TG-SC2A and VUB03TG-SC2B from VUB03KD-SC2, VUB03TG-SC3 from VUB03KD-SC3, **Figure 1A**) that were further cultured for 17 passages, i.e. till passage 29. These sublines all showed a significantly increased expression of the transgenic MSH2 protein compared to the *MSH2* knock-down lines which remained stable over time as demonstrated by immunocytochemistry (**Figure S8A**) and quantitative WB (**Figure S8B, D, Figure S9**). VUB03TG lines showed a 2- to 6-fold higher MSH2 expression compared to their MSH2KD counterparts with VUB03TG-SC2B (6-fold more than VUB03KD-SC2) showing the highest level of re-expression (**Figure S8B and D, Figure S9**). However, only VUB03TG-SC2B reached a level of MSH2 expression comparable to VUB03WT-SC1, while the other lines re-expressed MSH2 with a 2- to 3-fold lower expression when compared to the MSH2WT VUB03WT-SC1. MSH6 and MSH3 proteins equally showed a significant increase in protein level after MSH2 re-expression (**Figure S8D**).

We studied the CTG repeat size in these 4 lines at passage 29 and CpG methylation up- and downstream of the CTG repeat at passages 21, 25 and 29. In the presence of *MSH2* transgene expression, the CTG repeat size regained its unstable nature, with a bias towards expansions in all lines (**Figure 3, right panel**). Since the starting population at passage 12 only had a small variability in repeat size, we can infer that the observed increase in repeat size is due to the transgenic MSH2 and is not due to the selection of a cell with a larger expansion during cloning. The largest size increase was observed for VUB03TG-SC1 with an increase in the median size of 69 repeats over 17 passages, while the lowest was observed in VUB03TG-SC3 with an increase in the median size of only 4 repeats.

**Figure 3.**
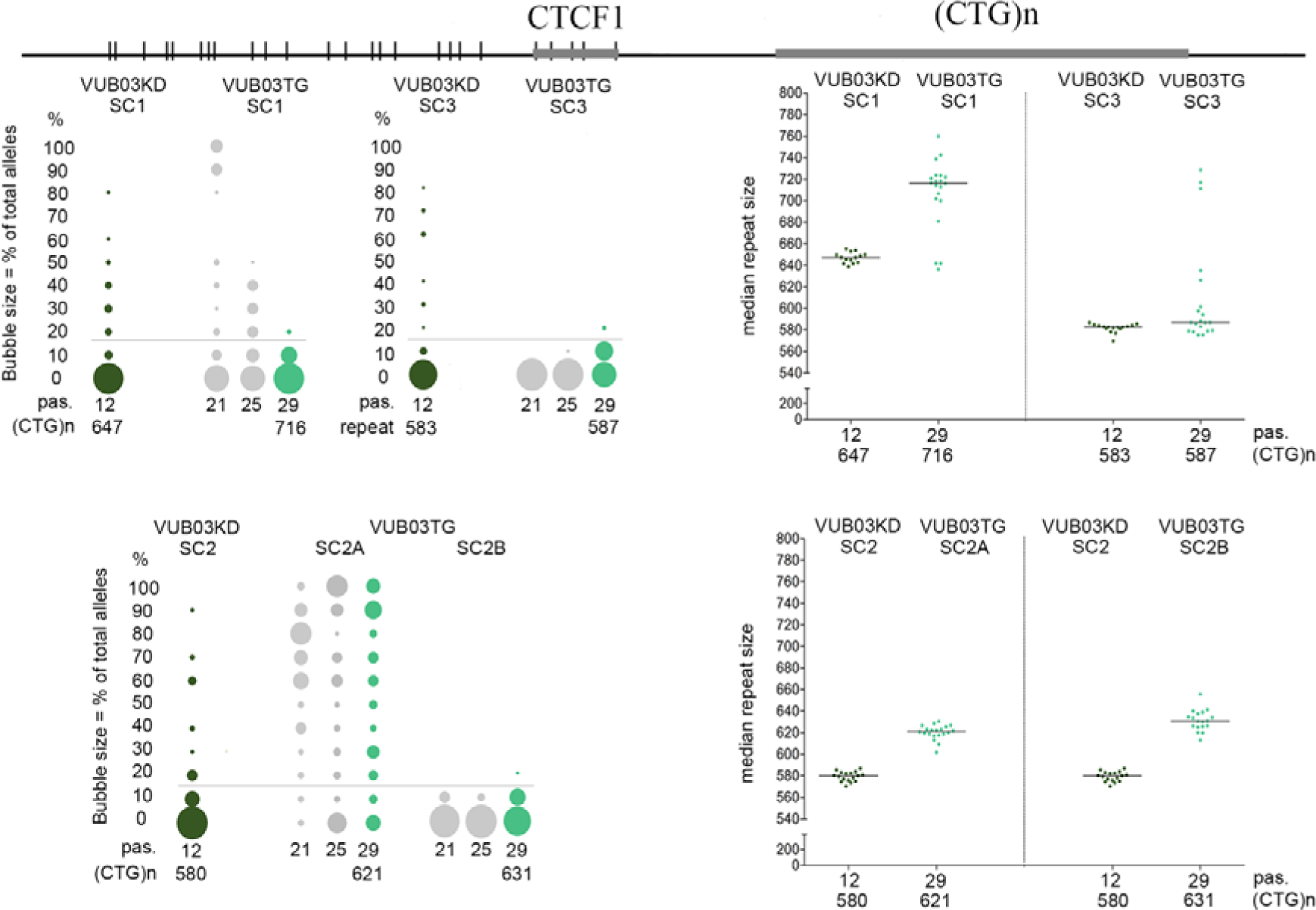
*MSH2* transgene expression re-initiates instability of the CTG repeat with a bias towards expansion but does not increase CpG methylation upstream of the CTG repeat. (A)The top of the graph schematically represents the *DMPK* CTG repeat region, with the 25 upstream CpG sites (vertical bars) and the CTCF1 site. The methylation levels are shown on the left and CTG length distributions are shown on the right per clonal line of VUB03-DM1 from passage 12 to passage 29. The X-axis shows the passage number and median CTG repeat size. Passage 12 (left data point) represents the upstream methylation of the bulk population just before *MSH2* transgene transduction. Passage 21, 25, 29 represent the upstream methylation pattern after *MSH2* transgene transduction. The right panel shows the median CTG repeat size for passage 12 and 29 for clonal lines of VUB03-DM1. Abbreviations: pas.: passage.

Lines VUB03TG-SC1, VUB03TG-SC2B and VUB03TG-SC3 show no significant difference in methylation pattern from their respective MSH2KD start point at passage 12 (Table **S5**). The two lines with the highest MSH2 re-expression, VUB03TG-SC1 and VUB03TG-SC2B, showed a marked increase in repeat length without re-establishing methylation. On the other hand, the re-expression in VUB03TG-SC3 (33% of VUB03WT-SC1, 2.4-fold more than VUB03KD-SC2) was low and this line only showed mild instability, with an increase of only 4 repeats in median size, combined with an absence of upstream methylation. Only 1 out of 4 lines (VUB03TG-SC2A) showed methylation upstream of the CTG repeat at passage 29 **(Figure 3, left panel)**, with 79% (**Table S6**) of the investigated epi-alleles being methylated. The methylation profile of VUB03TG-SC2A is significantly different from its original subline VUB03KD-SC2 at p12 (p= 0.023, **Table S5**). VUB03TG-SC2A also showed a lower increase in MSH2 re-expression (2-fold more than VUB03KD-SC2), however it did show an increase in repeat length combined with upstream CpG methylation. The methylation at the CpG region downstream of the CTG repeat (including the CTCF2 site) at the *DMPK* locus remains unchanged in all *MSH2* transgenic clonal lines (**Figure S10**). Given the discrepant results between the different lines, we analyzed the methylation status in the transduced bulk population, a few passages after the transduced passage 12 but before clonal expansion (**Figure S11**). All bulk MSH2TG lines show methylation similar to the methylation status in the MSH2KD lines at passage 12, i.e. VUB03KD-SC1_TG and VUB03KD-SC3_TG show low methylation as in VUB03KD-SC1 and VUB03KD-SC3, respectively, while VUB03KD-SC2_TG shows residual methylation similar to VUB03KD-SC2 (**Table S7**). It is therefore plausible that one clonal line obtained from VUB03KD-SC2_TG was methylated such as in VUB03TG-SC2A, originating from a cell carrying a methylated allele at the start of clonal expansion, while the other clone originated from a cell with a non-methylated allele resulting in VUB03TG-SC2B (**Figure S11**). This suggests that clonal selection is causing the variability in methylation at the onset of expansion of different TG sublines, and that this methylation status is not changed by the presence of MSH2 over time in culture.

In conclusion, cell lines with endogenous MSH2 expression show repeat instability over time in cell culture while knock down of MSH2 stabilized the repeat and even led to repeat contractions. Re-expression of MSH2 showed a re-initiation of repeat instability and repeat expansions. From this, we conclude that MSH2 plays a key role in repeat instability at the DMPK locus. The stabilization of the repeat upon *MSH2* knock-down coincided with the loss of methylation of the upstream CpG sites, establishing a connection between the MMR and the methylation status of this locus. Transgenic reintroduction of MSH2 did not re-establish CpG methylation in all tested lines.

## Discussion

In this study we performed *MSH2* knock-down experiments using CRISPR/Cas9 genome editing on two hESC lines VUB03-DM1 and VUB19-DM1 carrying a CTG expansion in the *DMPK* gene, and two non-DM1 hESC lines VUB02 and VUB06. We investigated the effect of the *MSH2* knock-down on both the CTG repeat dynamics and up and downstream CpG methylation on the CTCF sites at multiple time points during a cell culture lasting 20 passages.

We found that *MSH2* knock-down causes small contractions of the CTG tract and stabilization of the repeat, confirming similar findings reported in other model systems such as *Msh2-/-*DM1 mice^21^, an *in vitro* replication fork model^33^ and DM1 induced pluripotent stem cells (iPSCs)^23^. In our study, while loss of MSH2 halts further repeat expansion, the re-introduction of transgenic *MSH2* again leads to increased repeat size. Slean et al., (2016) were the first to propose a possible mechanism by which MutSβ protects against repeat contractions. They showed in an *in vitro* model that contraction intermediates are efficiently repaired leaving expansion-based slip-outs. In the absence of MutSβ, contraction-based DNA intermediates persist eventually leading to contractions of repeat size^33^. As such, not only a stabilization of the TNR, but also a contraction will occur.

The second finding of this study is that *MSH2* knock-down results in loss of CpG methylation upstream of the CTG repeat. This might suggest a causal association between MMR-mediated repeat instability and methylation upstream of the repeat. How this association between repeat size and methylation is established at the molecular level remains to be elucidated.

The causes and consequences of the presence of CpG methylation in the upstream region of the *DMPK* locus have been subject to much research, and several hypotheses have been tested, including a link between methylation and CTG repeat length and/or CTG repeat instability^7,19,26,28^. We previously demonstrated that hypermethylation of this region is strongly correlated with CDM1 and therefore with disease severity^19^, although the exact mechanism behind this correlation is still not clear. In addition, a strict correlation between methylation and CTG repeat size was not found^19^. It has also been suggested that an expanded CTG repeat itself is responsible for induction of methylation^29^. However, in a MSH2KD context, we observe methylation loss in the presence of repeat sizes falling within the range which was suggested by Brouwer *et al*. (2013) to induce methylation (DM300 and DMXL mice). This strengthens our hypothesis that MSH2 is a potential important factor for methylation of the CpG island upstream of the CTG repeat, including the CTCF1 site, at the *DMPK* region.

Transgenic *MSH2* expression in our knocked-down clonal lines results in increased CTG repeat instability but only resets methylation upstream of the CTG repeat in one of the four lines. This may indicate that continuous CTG repeat expansions or increased CTG repeat instability does not promote methylation in our DM1 hESC models, as proposed by Brouwer et al. (2013).

The connection between MSH2-induced repeat instability and upstream methylation may be indirect through a third partner. In an *in vitro* human model in which DNA double strand breaks (DSBs) were induced, it has been suggested that the MMR pathway recruits epigenetic modifiers such as the DNA methyl transferase DNMT1 at DNA damage sites in order to contribute to the DNA repair process^34–36^. After a successful repair, demethylase activity possibly restores the methylated site to its original unmethylated state^36^. In our MSH2WT hESC lines carrying a CTG expansion, CpG methylation flanking the CTG repeat is maintained and even increased over time in cell culture as in VUB19WT-SC1 and VUB19WT-SC2. This may be the result of repetitive DNA repair, which can lead to a permanent methylation signature, as seen in cancer cells^34^. This is consistent with the report of Yanovsky-Dagan *et al*. (2019) demonstrating demethylation upon CTG repeat excision in DM1 hESCs. However, they suggest that in undifferentiated hESCs, CpG methylation is re-established *de novo* after demethylation during DNA replication^37^. Our results might indicate that before a permanent methylated state is established, repetitive damage has to persist for a longer period of time.

Our results also show that this permanent methylated status is reversible exclusively upstream after eliminating a key component of the MMR pathway. Therefore, while the presence of MSH2 and a functional MMR may explain the maintenance of CpG methylation, other pathways must be involved in demethylation upon loss of MMR function. One possibility is passive demethylation during replication^38,39^. Another explanation would be that since the MMR component MSH2 is considerably reduced in our setting, another DNA repair mechanism might be at play. Goula et al. (2012) demonstrated that in a cellular environment with an active Base Excision Repair (BER) system such as the cerebellum, the TNR in Huntington’s disease is more stable and that long-patch BER (LP-BER) takes over in tissues with higher BER activity^40^. Although the TNR studied by these authors was not methylated, Franchini *et al*. (2014) showed that because of its processive nature demethylation through LP-BER may be a more efficient demethylation strategy than targeting single 5mC residues. It is therefore possible that demethylation occurs during LP-BER-mediated DNA repair, possibly with a preference of the upstream region due to the sensitivity of the LP-BER to the location of the lesion as shown by Goula *et al*. (2012). Re-methylation fails to occur because of the absence of MMR proteins. It is remarkable that only the upstream region is affected, as it was shown that the CTCF1 site at this location can actively bind CTCF protein and therefore play its role as insulator element, in which the downstream CTCF2 site is not involved^17,18^.

Re-expression of MSH2 into the VUB03TG lines caused the repeat to become unstable again, although to a much smaller extent than in VUB03WT lines: while WT lines easily gained 200 repeats over 16 passages, this was only about 50 repeats for the TG lines. Concurrently, upstream CpG methylation was not reintroduced and was only present in one clonal line presumably originating from a cell with a methylated allele. Possibly, the reintroduction of MSH2 which was not to the same level as in the VUB03WT lines was nevertheless high enough to cause mild instability, but not methylation as seen over time in VUB19WTSC1 and VUB19WTSC2. These data might suggest that the involvement of MSH2 in upstream CpG methylation is dose-dependent.

In conclusion, our data not only confirm the role of the MMR in repeat instability, but also suggest a connection between the MMR pathway and the introduction and maintenance of CpG methylation upstream of the CTG repeat in DM1. MMR-mediated repair could possibly initiate aberrant DNA methylation which is then maintained by a constant recruitment of the methylation machinery during a continued repair of intermediate DNA structures. In contrast, the mechanism behind the observed methylation loss could be a passive demethylation upon loss of the MMR pathway, or might be induced by LP-BER leading to rapid demethylation. Repeat instability caused by the MMR pathway and CpG methylation at the repeat have been linked to disease severity. Our data provide a starting point to understand a possible crosstalk between CTG repeat instability and flanking CpG methylation and could contribute to a better understanding of the mechanisms underlying the DM1 disease severity.

## Materials and Methods

### Human embryonic stem cell culture

Human ESCs were derived from pre-implantation embryos as described in ^8,9,41^ after obtaining the patient’s consent and with the permission of the Commission for Medical Ethics of the UZ Brussel and the Federal Commission for Research on Embryos. All cell lines are registered in the EU hPSC registry, https://hpscreg.eu/. All hESC lines were cultured on 10 µg/mL recombinant laminin-521 (Biolaminin 521 LN; Biolamina) coated dishes in NutriStem^®^ hESC XF medium (Biological Industries), supplemented with 100 U/mL penicillin/streptomycin (Pen/Strep; ThermoFisher). Cell lines were cultured at 37°C in 5% CO_2_ and atmospheric O_2_ conditions. Medium was changed daily. Cell passaging was performed weekly using 1x TrypLE^™^ Express (ThermoFisher) and cells were replated in a 1:10 to 1:50 ratio depending on the growth speed of the cell line.

### Analysis of CTG repeat instability: long CTG repeat analysis by PacBio sequencing

In order to amplify only one to five DNA molecules per reaction, we used a small pool PCR with an input of 20 to 50 pg as described in Seriola et al. (2011). For each cell DNA sample, 20 PCR reactions with low input template DNA were analysed to establish the distribution of the repeat sizes in each sample ^8,9,19^. Repeats were amplified with high fidelity using the LongAmp Taq polymerase (New England Biolabs). Twenty to 50pg of DNA was amplified in a 25 µL reaction mix containing 2.5 units LongAmp Taq DNA polymerase, 1x LongAmp buffer (New England Biolabs), 0.2 mM dNTPs (Illustra DNA polymerization mix, GE Healthcare) and 0.4 µM of primers DM101 and DM102 (Integrated DNA Technologies)^2^ and 2.5% dimethyl sulphoxide (DMSO). Primer sequences are listed in **Materials and Methods S1**. Amplification conditions were as follows: 4 min of initial denaturation at 94°C, 35 cycles of 30s denaturation at 94°C, 8min annealing and extension at 65°C and a final extension step at 65°C for 10min. The LongAmp amplicons, spanning the repeat, were prepared for sequencing as described in PacBio’s guide for Preparing SMRTbell^™^ Libraries using PacBio^®^ Barcoded Adapters for Multiplex SMRT^®^ Sequencing. This protocol allows to pool 2 samples which each consist of 20 PCR products with a different barcode in one library. Before exonuclease treatment, 500 ng of PUC19 plasmid was added to avoid degradation of intact SMRTbells. Libraries were purified with Ampure beads multiple times, according to PacBio’s instructions, in order to eliminate small repeat size fragments and to favour the sequencing of the longer alleles. Each library was sequenced on a single SMRT cell by a PacBio RS II or Sequel using the DNA/Polymerase binding Kit P6 v2 (Pacific Biosciences) for a 360 min’ movie. We used PacBio’s DNA Sequencing Reagent Kit 4.0 v2 for all runs. Therefore, demultiplexed circular consensus (CCS) reads were generated with the RS_ReadsOfInsert.1 protocol from PacBio’s SMRT portal (v2.3.0) or with ccs and lima software from SMRTLink (v6.0.0) with a minimum of 1 full pass, a minimum predicted accuracy of 90% and demultiplexing with symmetric barcodes. Next, each PCR product was aligned to the *DMPK* CTG repeat using BWA-SW v0.7.10^42^ against the human reference genome hg19 downloaded from UCSC^43^, followed by conversion of SAM to BAM by Samtools v1.3.1^44^. To finally convert to BED format and select the on-target CCS reads BEDtools v2.20.1 was used ^45^. For each CCS read spanning the CTG repeat, the number of repeat units was determined by measuring the distance between two unique regions flanking the CTG repeat followed by detecting the most abundantly present repeat size in each PCR product, here represented by the median.

### *MSH2* knock-down using CRISPR/Cas9 and CRISPR/Cas9nickases

Two different genome editing CRISPR/Cas9 strategies, CRISPR/Cas9 and CRISPR/Cas9nickases were used to create *MSH2* knock-down cell lines. CRISPR/Cas9 uses the wild type Cas9 enzyme and creates a double stranded break on a specific location in the DNA, while single stranded breaks are obtained by means of a point mutated Cas9 enzyme called Cas9nickase (Cas9Nick) (**Materials and Methods S2**). Both methods showed equal genome editing efficiencies and were able to induce random mutations in both alleles by non-homologous end joining repair, and subsequently knocked-down *MSH2*.

CRISPR/Cas9 and Cas9Nick sgRNA’s (**Materials and Methods S1**) were designed using the online CRISPR design tool (https://zlab.bio/guide-design-resources) ^46^. sgRNA Cas TOP MSH2 ex3 and sgRNA Cas BOTTOM MSH2 ex3 are the oligonucleotides used for CRISPR/Cas9, the forward guide used for Cas9Nick genome editing consists of the oligonucleotides: sgRNACas9n forward TOP and BOTTOM and the reverse guide of sgRNACas9n reverse TOP and BOTTOM. The appropriate sgRNA oligonucleotides were individually annealed and cloned in a Cas9 or Cas9Nick containing vector: pSpCas9(BB)-2A-Puro (PX459) (plasmid#48139, Addgene) and pSpCas9n(BB)-2A-Puro (PX461) (plasmid#48141, Addgene) respectively, according to Ran *et al*. 2013^46,47^. Vectors were generated in competent Stbl3^™^ *E. coli* cells with the One shot^®^ Stbl3^™^ chemically competent *E. coli* kit (Invitrogen). The transfection was performed according to the manufacturer’s guidelines and vector assembly was tested by Sanger sequencing.

Human ESC lines were plated in high density one day before transfection in order to reach 50% confluency at the time of transfection. Cells were transfected using Lipofectamine^®^ 3000 (Invitrogen), according to manufactures guidelines. Twelve hours after transfection, 0.8 to 2 µg/mL puromycin (Sigma-Aldrich) selection, depending on the cell line, was performed for twenty-four hours. Non-transfected cells were used as control for a successful puromycin selection. After puromycin selection, cells were grown until colonies were visible and subsequently passaged at low densities on Biolaminin 521 LN® (Biolamina) coated 24 well plates supplemented with 1.5 µg/mL E-Cadherin (R&D systems) to obtain single cell colonies. Subsequently single cell colonies were picked manually and expanded for downstream experiments. Single cell colonies were obtained from wild type hESC lines to serve as a control.

### DNA, RNA extraction and cDNA conversion

DNA extraction was obtained from pelleted cells, according to the DNeasy Blood & Tissue kit and DNeasy Micro kit (QIAGEN). Total RNA was extracted from pelleted cells using the Rneasy Mini kit (QIAGEN) following manufacturer’s guidelines for animal spin cells. DNA and RNA concentrations were measured using the Nanodrop and the Qubit using the Qubit^®^ dsDNA Broad Range and RNA Assay kit (Life Technologies). DNA was stored at 4°C, RNA at −80°C. RNA was reverse-transcribed to cDNA using the First-Strand cDNA Synthesis Kit (GE Healthcare) following manufacturer’s guidelines and was stored at −20°C.

### *MSH2* gene editing validation with PCR and Sanger sequencing and MiSeq

Exon 3 of the MSH2 gene was PCR amplified using the Expand High Fidelity PCR kit (Roche) with the following conditions: initial denaturation at 94°C for 2min, followed by 30 cycles of denaturation at 94°C (15s), annealing at 54°C (30s) and extension at 72°C (45s). The latter step is expanded with a time increase of 5s per cycle starting from cycle 11. This is followed by an extension step at 72°C for 7min. *MSH2* exon3 primers can be found in **Materials and Methods S1**. PCR products were purified using the High Pure PCR Product Purification Kit (Roche) following manufacturer’s guidelines. Subsequently, a BigDye reaction was performed on the purified product, according to the BigDye^®^ Terminator v3.1 Cycle Sequencing Kit (Applied Biosystems). The PCR program contained the following steps: initial denaturation at 96°C for 1min, followed by 25 cycles of denaturation at 96°C (10s), annealing at 50°C (5s) and extension at 60°C (4min). Samples were purified, run on an ABI 3130XL automatic sequencer (Applied Biosystems) and analyzed using Applied Biosystems Sequence Scanner v1.0 software. A more precise validation of the introduced mutation in the *MSH2* gene was obtained after MiSeq. MSH2 PCR products were purified using 1,8x AMPure XP bead cleanup (Beckman Coulter Life Sciences, IN, USA) and eluted in 30 μl of nuclease free water. A total of 80 ng of purified amplicon was inputted in the KAPA Hyper Prep (Roche Sequencing, CA, USA) library preparation according to manufacturer’s recommendations, with four exceptions: (1) the volumes were reduced with a factor 2, (2) Unique Dual Indexed (UDI) adapters of own design (Integrated DNA Technologies, Coralville, IA, USA) were used at a concentration of 15 μM, (3) a 1x post ligation bead cleanup with AMPure XP beads (Beckman Coulter Life Sciences, IN, USA) was performed and (4) a total of 6 PCR cycles were applied to obtain sufficient library DNA. All libraries were sequenced on a MiSeq instrument (Illumina Inc., CA, USA) using the MiSeq v2 reagent kit, generating 2×250bp reads. For this, libraries were denatured and diluted to a final concentration of 10 pM and a 15% PhiX control library was included in the sequencing run according to manufacturer’s instructions. Base called reads in the fastq format were aligned against the human reference genome hg19 assembly ^48^ using bwa mem ^49^. The generated files in the .bam format were sorted and indexed with samtools ^44^. Low-frequency somatic variants were called using Pisces ^50^ and the generated .vcf files were annotated with Alamut batch 1.9 (standalone) / Qt 5.3.2 (Interactive Biosoftware, Rouen, France).

### Off-target analysis of CRISPR/Cas9

Eight potential off-target sites were computationally identified (https://zlab.bio/guide-design-resources) and analyzed for possible mutations caused by the wild type Cas9 enzyme. The computationally generated off-target sites were amplified with primer-specific annealing conditions (**Materials and methods S1**) and subsequently sequenced by Sanger sequencing. Briefly, 10 ng of DNA was amplified using AmpliTaq on a thermocycler (Life Technologies). The total reaction volume of 25 µL contained 1.25 units AmpliTaq DNA polymerase (Applied Biosystems), 25 mM MgCl_2_ (Applied Biosystems), 1x PCR buffer (Applied Biosystems), 0.2 mM dNTPs (Illustra DNA polymerization mix, GE Healthcare) and 0.4 µM primers (Integrated DNA Technologies, Supplementary Table 1). The following PCR program was used: initial denaturation at 95°C for 5 min, followed by cycles of denaturation at 95°C (30 s), annealing at primer-specific temperatures (30 s) and extension at 72°C (30 s). The amount of cycles differed dependent on the gene that was amplified (Supplementary Table 1). A final extension at 72°C for 5 min was used. The amplification of the C5 region was performed with a nested PCR following the conditions mentioned in **Materials and methods S1**.

### RT-qPCR to quantify mRNA expression levels

mRNA expression of *MSH2, MSH3* and *MSH6* (Hs00179887_m1; Hs00989003_m1 and Hs00264721_m1 respectively) in knock-down lines was compared to non-transfected cell lines using real-time qPCR. The 20 µL reaction mix contained 40 ng cDNA, 10 µL TaqMan^®^ Fast Advanced Master Mix (Applied Biosystems) and either 1 µL TaqMan assay (ThermoFisher Sientific) for *MSH2, MSH3, MSH6* and *GUSB* (Hs99999908_m1) or 1.8 µM primer mix (IDT) and 250 nM probes (ThermoFisher Scientific) for *GAPDH* and *UBC. GUSB, GAPDH* and *UBC* were used as endogenous controls. Primer and probe sequences can be found in **Materials and Methods S1**. Samples were run on a ViiA7 Real-Time PCR system (ThermoFisher Scientific) and analysed with VIIA7 software v1.2 (ThermoFisher Scientific). Cycling conditions were as follows: 95°C for 20s followed by 40 cycles of 95°C for 1s and 60°C for 20s.

### Immunocytochemistry for MSH2

Cells were fixed with 4% paraformaldehyde (PFA) for 10 min at room temperature, washed twice with PBS and permeabilized using 0.3% Triton-X (Sigma-Aldrich) for 10min ^51^. Briefly, after permeabilization cells were blocked with 3% BSA, 0.1% Tween in PBS for 30min. Subsequently, cells were incubated with mouse anti-human MSH2 primary antibody (1:500) (ab52266, Abcam) diluted in 0.1% BSA, 0.1% Tween and PBS for 1h. After two washes, cells were incubated with the first antibody (1/50, DSHB) diluted in 0.1% BSA, 0.1% Tween and PBS for 45min and subsequently stained with the secondary antibody (1/500) diluted in 0.1% BSA, 0.1% Tween and PBS for 30min. Nuclear staining was performed with Hoechst 33342 (Thermo Fischer Scientific). Images were taken by confocal microscopy using LSM800 (Zeiss). Each blot was repeated twice.

### Bisulfite treatment and Massive Parallel Sequencing or Sanger sequencing

Bisulfite treated massive parallel sequencing was performed as described in ^19^. Briefly, the Imprint DNA Modification Kit (Sigma Aldrich) was used for bisulfite treatment on 200ng DNA. Bisulfite-treated DNA was amplified using primers in **Materials and Methods S1** for regions upstream containing 25 CpG sites and including the CTCF1 and downstream (11 CpG sites, CTCF2) of the CTG repeat, using the Jumpstart Taq DNA Polymerase Kit (Sigma Aldrich). The first and second round PCR conditions were adapted from Barbé *et al*., 2017^19^. First round PCR primers (**Materials and Methods S1**) are indicated by ‘1’ at the end of the target name, second round primers (**Materials and Methods S1**) are indicated by Miseq at the end of the target name. Libraries were made as described in ^19^ and subsequently loaded on the MiSeq Reagent Nano Kit v2 (500 cycles) according to manufacturer’s instructions and sequenced at 2×250 bp (Illumina). During data analysis, 100 sequences for each region, with a length more than 150bp, were processed using a homemade script. BiQ Analyzer aligns the sequences to the reference using Crustal W alignment. Sequences with less than 90% bisulfite conversion rate and/or less than 80% sequence identity with the reference sequence were discarded. Methylation status, percentage of methylated CpG sites of the total number (25 in the upstream region, 11 in the downstream region) of potentially methylated sites, was calculated for each of these 100 random selected sequences individually. These 100 sequences per regions were binned according to the calculated methylation percentage: no methylation as seen on wild type alleles (under 10%) and methylation (10-100%).

Non-DM1 hESC lines were Sanger sequenced. The PCR product was cycle-sequenced using BigDye chemistry 3.1 according to the manufacturer’s guidelines (Life Technologies). Subsequently, amplicons were run on an ABI 3130XL automatic sequencer (Applied Biosystems). Results were analyzed using Applied Biosystems Sequence Scanner v1.0. A methylated CpG site shows two peaks, one for a C (methylated) and one for a T (unmethylated). An unmethylated CpG site only shows one peak, one for a T (unmethylated).

### Western blot for protein expression

Proteins were extracted from a cell pellet by resuspension in a non-denaturing lysis buffer containing 20mM Tris HCl pH8, 137 mM NaCl, 1% NP-40, 2mM EDTA and cOmplete^™^ Protease inhibitor (Sigma Aldrich). After 1-hour incubation on ice, cells were centrifuged for 10 min at 4°C at 21000g. The supernatant was collected and protein concentrations were measured with the Qubit Protein Assay kit (Life Technologies). Protein samples were diluted in 4x sample buffer (4x Laemmli buffer + β-mercaptoethanol), obtaining a protein concentration between 10-30 µg in a total volume of 25µL. Before loading, diluted samples were boiled for 5min at 95 °C and subsequently separated on a 4-15% Criterion^™^ TGX Stain-Free^™^ Protein Gel (Bio-Rad Laboratories) and ran in a Criterion^™^ Cell geltank (Bio-Rad Laboratories) at 200V for 45min in TGS running buffer (Bio-Rad Laboratories). Proteins were transferred on a nitrocellulose membrane using a semi-dry blotting technique at 25V for 7min. Subsequently nitrocellulose membranes were incubated with target primary antibodies (**Materials and Methods supplement 1**) overnight. The next day, membranes were washed three times with PBS + 0.1% Tween20 for 5min each wash and later incubated with fluorescently labelled secondary antibodies (**Materials and Methods supplement 1**) for 60min in the dark. After three additional washing steps, bands were visualized using the Odyssey Infrared Imager (LI-COR Biosciences). Relative protein amounts were calculated relative to the housekeeping protein Actin and a control using Image Studio software and statistically evaluated with a t-test (p<0,05).

### *MSH2* transgene expression

The *MSH2* gene was reintroduced in the *MSH2* knock-down hESC lines using lentiviral vectors which were made by Origene Technologies GmbH. They re-cloned the open reading frame of RC205848 in PS100106 and produced lentiviral particles carrying the *MSH2* construct. Cells were transduced when they reached 50% confluency. Cells were subsequently selected using 0.8 µg/mL puromycin (Sigma-Aldrich). Depending on the survival rate of the non-transduced controls, cells were selected twice. After selection recovery, cells were plated as single cells in order to obtain *MSH2*+/+ clonal lines as described in the section ‘*MSH2* knock-down using CRISPR/Cas9 and CRISPR/Cas9nickases’. Subsequently cells were cultured over multiple passages.

### Statistics

Our PacBio data do not show a normal distribution and are therefore unfit for the use in parametrical statistical tests. In addition, our data within a clonal line including passage 4, 12 and 20, are ordered in a specific manner. Therefore we used the Jonckheere Terpstra Test which is based on comparing medians to study differences in median repeat size across cell lines and conditions^52^. A result of p<0.05 in the Jonckheere-Terpstra test indicates that our data follows a specific trend. For the methylation data, a Kolmogorov-Smirnov test was performed to compare the pair-wise distribution of two samples. A statistical significance (p<0.05) indicates that data distributions are not the same and were considered as a statistically different methylation profile.

## Acknowledgements

We thank Prof. dr. Christopher Pearson and dr. Stella Lanni (Toronto, Canada) for their support and constructive criticisms. We also thank Prof. dr. Kurt Barbé for his help with the statistical analysis. We acknowledge the technical support by the team of BRIGHTCore for the Miseq analysis and Sanger sequencing. We also acknowledge the Genomics Core at UZ Leuven for their technical support for PacBio sequencing. This work was supported by the Methusalem grant in name of Karen Sermon and FWO grant G.0223.15N and by the Hercules foundation (ZW11-14).

## Conflict of interest statement

None Declared

## Funding

This work was supported by the Methusalem grant in name of Karen Sermon and FWO grant G.0223.15N and by the Hercules foundation (ZW11-14).

## Abbreviations

DM1: myotonic dystrophy type 1
CTCF1: CCCTC-binding factor site 1
CTCF2: CCCTC-binding factor site 2
MMR: mismatch repair pathway
TNR: trinucleotide repeat
hiPSCs: human induced pluripotent stem cells
hESCs: human embryonic stem cells
OPLs: osteogenic progenitor-like-cells
KD: knock-down
WT: wild type
SC: single cell clonal line
pas: Passage
TG: transgenic expression
DSBs: DNA double strand breaks
LP-BER: long patch base excision repair

## Notes

### Competing Interest Statement

The authors have declared no competing interest.

## References

1. Udd, B. & Krahe, R. The myotonic dystrophies: Molecular, clinical, and therapeutic challenges. Lancet Neurol. 11,891–905 (2012).

2. Brook, J. D. et al. Molecular basis of myotonic dystrophy: expansion of a trinucleotide (CTG) repeat at the 3’ end of a transcript encoding a protein kinase family member [published erratum appears in Cell 1992 Apr 17;69(2):385]. Cell 68,799–808 (1992).

3. Lanni, S. & Pearson, C. E. Molecular genetics of congenital myotonic dystrophy. Neurobiol. Dis. 104533 (2019). doi: 10.1016/j.nbd.2019.104533

4. Mirkin, S. M. Expandable DNA repeats and human disease. Nature 447,932–40 (2007).

5. Schmidt, M. H. M. & Pearson, C. E. Disease-associated repeat instability and mismatch repair. DNA Repair (Amst). 38,117–126 (2016).

6. Pearson, C. E. & Sinden, R. R. Alternative structures in duplex DNA formed within the trinucleotide repeats of the myotonic dystrophy and fragile X loci. Biochemistry 35,5041–5053 (1996).

7. Lopez-Castel, A. et al. Expanded CTG repeat demarcates a boundary for abnormal CpG methylation in myotonic dystrophy patient tissues. Hum. Mol. Genet. 20,1–15 (2011).

8. De Temmerman, N. et al. CTG repeat instability in a human embryonic stem cell line carrying the myotonic dystrophy type 1 mutation. Mol. Hum. Reprod. 14,405–412 (2008).

9. Seriola, A. et al. Huntington’s and myotonic dystrophy hESCs: down-regulated trinucleotide repeat instability and mismatch repair machinery expression upon differentiation. Hum. Mol. Genet. 20,176–85 (2011).

10. Wohrle, D. et al. Heterogeneity of DM kinase repeat expansion in different fetal tisseus and further expansion during cell proliferation in vitro: evidence for a causal involvment of methyl-directed DNA mismatch repair in triplet repeat stability. Hum. Mol. Genet. 4, (1995).

11. Wong, L. J. C., Ashizawa, T., Monckton, D. G., Caskey, C. T. & Richards, C. S. Somatic heterogeneity of the CTG repeat in myotonic-dystrophy is age and size-dependent. Am. J. Hum. Genet. 56,114–122 (1995).

12. Morales, F. et al. Somatic instability of the expanded CTG triplet repeat in myotonic dystrophy type 1 is a heritable quantitative trait and modifier of disease severity. Hum. Mol. Genet. 21,3558–3567 (2012).

13. Brook, J. D. et al. Molecular basis of myotonic dystrophy: Expansion of a trinucleotide (CTG) repeat at the 3′ end of a transcript encoding a protein kinase family member. Cell 68,799–808 (1992).

14. De Temmerman, N. et al. Intergenerational instability of the expanded CTG repeat in the DMPK gene: studies in human gametes and preimplantation embryos. Am. J. Hum. Genet. 75,325–329 (2004).

15. Howeler, C. J., Busch, H., Geraedts, J., Niermeijer, M. & Staal, A. Anticipation in myotonic dystrohy: Fact or fiction. Brain 112,779–797 (1989).

16. Johnson, N. E. et al. Disease burden and functional outcomes in congenital myotonic dystrophy: A cross-sectional study. Neurology 87,160–167 (2016).

17. Cho, D. H. et al. Antisense transcription and heterochromatin at the DM1 CTG repeats are constrained by CTCF. Mol. Cell 20,483–489 (2005).

18. Filippova, G. N. et al. CTCF-binding sites flank CTG/CAG repeats and form a methylation-sensitive insulator at the DM1 locus. Nat. Genet. 28,335–343 (2001).

19. Barbé, L. et al. CpG Methylation, a Parent-of-Origin Effect for Maternal-Biased Transmission of Congenital Myotonic Dystrophy. Am. J. Hum. Genet. 100,488–505 (2017).

20. Usdin, K., House, N. C. M. & Freudenreich, C. H. Repeat instability during DNA repair: Insights from model systems. Crit. Rev. Biochem. Mol. Biol. 50,142–167 (2015).

21. Savouret, C. et al. CTG repeat instability and size variation timing in DNA repair-deficient mice. EMBO J. 22,2264–2273 (2003).

22. Tomé, S. et al. MSH2 ATPase domain mutation affects CTG-CAG repeat instability in transgenic mice. PLoS Genet. 5, (2009).

23. Du, J., Campau, E., Soragni, E., Jespersen, C. & Gottesfeld, J. M. Length-dependent CTG.CAG triplet-repeat expansion in myotonic dystrophy patient-derived induced pluripotent stem cells. Hum. Mol. Genet. 22,5276–5287 (2013).

24. van den Broek, W. J. A. A. et al. Somatic expansion behaviour of the (CTG)n repeat in myotonic dystrophy knock-in mice is differentially affected by Msh3 and Msh6 mismatch-repair proteins. Hum. Mol. Genet. 11,191–8 (2002).

25. Spits, C., Seneca, S., Hilven, P., Liebaers, I. & Sermon, K. Methylation of the CpG sites in the myotonic dystrophy locus does not correlate with CTG expansion size or with the congenital form of the disease. J. Med. Genet. 47,700–703 (2010).

26. Nakamori, M. et al. Aberrant Myokine Signaling in Congenital Myotonic Dystrophy. Cell Rep. 21,1240–1252 (2017).

27. Steinbach, P., Gläser, D., Vogel, W., Wolf, M. & Schwemmle, S. The DMPK gene of severely affected myotonic dystrophy patients is hypermethylated proximal to the largely expanded CTG repeat. Am. J. Hum. Genet. 62,278–285 (1998).

28. Yanovsky-Dagan, S. et al. Uncovering the role of hypermethylation by CTG expansion in myotonic dystrophy type I using mutant humand embryonic stem cells. Stem Cell Reports 5,1–11 (2015).

29. Brouwer, J. R., Huguet, A., Nicole, A., Munnich, A. & Gourdon, G. Transcriptionally repressive chromatin remodelling and CpG methylation in the presence of expanded CTG-repeats at the DM1 locus. J. Nucleic Acids 2013, (2013).

30. Acharya, S. et al. hMSH2 forms specific mispair-binding complexes with hMSH3 and hMSH6. Proc. Natl. Acad. Sci. U. S. A. 93,13629–34 (1996).

31. Chang, D. K., Ricciardiello, L., Goel, A., Chang, C. L. & Boland, C. R. Steady-state regulation of the human DNA mismatch repair system. J. Biol. Chem. 275,18424–18431 (2000).

32. Ardui, S. et al. Detecting AGG interruptions in females with a FMR1 premutation by long-read single-molecule sequencing: A 1 year clinical experience. Front. Genet. 9,1–6 (2018).

33. Slean, M. M. et al. Absence of MutSBeta leads to the formation of slipped-DNA for CTG/CAG contractions at primate replication forks. DNA Repair (Amst). 42,107–118 (2016).

34. Ding, N. et al. Mismatch repair proteins recruit DNA methyltransferase 1 to sites of oxidative DNA damage. J. Mol. Cell Biol. 0,1–11 (2015).

35. Ding, N., Miller, S. A., Savant, S. S. & O’Hagan, H. M. JAK2 regulates mismatch repair protein-mediated epigenetic alterations in response to oxidative damage. Environ. Mol. Mutagen. 60,308–319 (2019).

36. Ding, N., Maiuri, A. R. & O’Hagan, H. M. The emerging role of epigenetic modifiers in repair of DNA damage associated with chronic inflammatory diseases. Mutat. Res. - Rev. Mutat. Res. 780,69–81 (2019).

37. Yanovsky-Dagan, S. et al. Deletion of the CTG Expansion in Myotonic Dystrophy Type 1 Reverses DMPK Aberrant Methylation in Human Embryonic Stem Cells but not Affected Myoblasts. bioRxiv (2019). doi: 10.1101/631457

38. Grin, I. & Ishchenko, A. A. An interplay of the base excision repair and mismatch repair pathways in active DNA demethylation. Nucleic Acids Res. 44,3713–3727 (2016).

39. Franchini, D. M. et al. Processive DNA demethylation via dna deaminase-induced lesion resolution. PLoS One 9, (2014).

40. Goula, A.-V. et al. Nucleotide sequence, DNA damage location and protein stoichiometry influence base excision repair outcome at CAG/ CTG repeats. Biochemistry 51,3919–3932 (2012).

41. Mateizel, I. et al. Derivation of human embryonic stem cell lines from embryos obtained after IVF and after PGD for monogenic disorders. Hum. Reprod. 21,503–511 (2006).

42. Li, H. & Durbin, R. Fast and accurate short read alignment with Burrows-Wheeler transform. Bioinformatics 25,1754–1760 (2009).

43. Karolchik, D. et al. The UCSC Table Browser data retrieval tool. Nucleic Acids Res. 32,493–496 (2004).

44. Li, H. et al. The Sequence Alignment/Map format and SAMtools. Bioinformatics 25,2078–2079 (2009).

45. Quinlan, A. R. & Hall, I. M. BEDTools: A flexible suite of utilities for comparing genomic features. Bioinformatics 26,841–842 (2010).

46. Ran, F. A. et al. Genome engineering using the CRISPR-Cas9 system. Nat. Protoc. 8,2281–2308 (2013).

47. Ran, F. A. et al. Double nicking by RNA-guided CRISPR cas9 for enhanced genome editing specificity. Cell 154,1380–1389 (2013).

48. Lander, E. S.. L. et al. Initial sequencing and analysis of the human genome. Nature 409,860–921 (2001).

49. Li, H. Aligning sequence reads, clone sequences and assembly contigs with BWA-MEM. 00, 1–3 (2013).

50. Dunn, T. et al. Pisces: An accurate and versatile variant caller for somatic and germline next-generation sequencing data. Bioinformatics 35,1579–1581 (2019).

51. van der Wal, E. et al. GAA Deficiency in Pompe Disease Is Alleviated by Exon Inclusion in iPSC-Derived Skeletal Muscle Cells. Mol. Ther. - Nucleic Acids 7,101–115 (2017).

52. Ali, A. et al. Non-Parametric Test for Ordered Medians: The Jonckheere Terpstra Test. Int. J. Stat. Med. Res. 4,217–233 (2015).

