## Supplementary data for "*MSH2* knock-down shows CTG repeat stability and concomitant upstream demethylation at the *DMPK* locus in myotonic dystrophy type 1 human embryonic stem cells"

**Table S1.** Overview of the oligos and antibodies used. Listed are primers for LongAmp PCR spanning the repeat and for identifying mutations in *MSH2* exon3, the gRNAs for CRISPR/Cas9 and CRISPR/Cas9Nick, the primers used for identifying off-targets, the primers used for RT-qPCR endogenous controls and the primers for massive parallel sequencing of the CpG regions up- and downstream of the CTG. ‘F’ indicates the forward primers, ‘R’ the reverse primers. Number ‘1’ following ‘F’ or ‘R’ indicate the first round of the nested PCR. For the second PCR round, primers with Miseq at the end of the target name were used. An overview of antibodies used in immunocytochemistry is shown.

| target name | forward primer | reverse primer |
|---|---|---|
| DM101/102 | 5'-CTTCCCAGGCCTGCAGTTTGCCCATCCA-3' | 5'-GAACGGGGCTCGAAGGGTCCTTGT-3' |
| *MSH2* exon 3 | 5'-CAAAAAGGAAAAATCCAAACTCTATG-3' | 5'-CCTTTGGTCCAATCTGGATG-3' |

| sgRNA's name | sequence TOP strand | sequence BOTTOM strand |
|---|---|---|
| sgRNACas9 | 5'-CACCGGTTAAAATGTCCGCAGTTGA-3' | 5'-AAACTCAACTGCGGACATTTTAACC-3' |
| forward sgRNACas9n | 5'-CACCGGGTATGTGGATTCCATACAG-3' | 5'-CACCGCTGTCTCTGGCCATCAACTG-3' |
| reverse sgRNACas9n | 5'-AAACCTGTATGGAATCCACATACCC-3' | 5'-AAACCAGTTGATGGCCAGAGACAGC-3' |

| off-target name | forward primer | reverse primer | PCR condition |
|---|---|---|---|
| *LINC00882* | 5'-GCAGGAGGATCTATGGGTGA-3' | 5'-TGGAGAAAGGGTCAGGTCAA-3' | an: 62°C, 5% DMSO, 30 cycles |
| *F9* | 5'-CCGGGCATTCTAAGCAGTTT-3' | 5'-GAGGGAAACTTTGAACCATGAGT-3' | an: 60°C, 30 cycles |
| *PGAP3* | 5'-GATTCCCTCATCCTGCTCCA-3' | 5'-TGAGTGCATAGGTGACAGGG-3' | an: 62°C, 5% DMSO, 30 cycles |
| *ACACA* | 5'-CATGGCAACCTCTGGATTGG-3' | 5'-TTGCACTACCTTGGCTGGAT-3' | an: 60°C, 30 cycles |
| *UTRN* | 5'-CAATGCCTACTTTGTTTCCACA-3' | 5'-ACTGTAAAGCAAAATCAAGGTGG-3' | an: 60°C, 30 cycles |
| *SON* | 5'-CAATGAATCTGTCACCAAGTTC-3' | 5'-TGGGTTCCTTATTGCCTTCTT-3' | an: 60°C, 30 cycles |
| *FAM63A* | 5'-TGAGGAGACGGTCTGAGGTA-3' | 5'-TCACCTGAGTCATTCCCTGG-3' | an: 60°C, 30 cycles |
| *C5* | <u>1st PCR</u><br>5'-CTCTACTTTCTGGCGCACAC-3'<br><u>2nd PCR</u><br>5'-AGGACTTTGTGCCCTGATGA-3' | <u>1st PCR</u><br>5'-AATCCCAGCTACTCAGGAGG-3'<br><u>2nd PCR</u><br>5'-GGAGAATTGCTTGAATCCGGG-3' | <u>1st PCR</u><br>an: 62°C, 5% DMSO, 25cycles<br><u>2nd PCR</u><br>an: 62°C, 5% DMSO, 18 cycles |

| target name | forward primer | reverse primer | probe |
|---|---|---|---|
| *GAPDH* | 5'-ATGGAAATCCCATCACCATCTT-3' | 5'-CGCCCCACTTGATTTTGG-3' | 6-FAM-CAGCAGCGAGATCC-MGB |
| *UBC* | 5'-CGCAGCCGGGATTTG-3' | 5'-TCAAGTGACGATCACAGCGA-3' | 6-FAM-TCGCAGTTCTTGTTTGTG-MGB |

| target name | primer sequence |
|---|---|
| CTCF1 F1 | 5'-TGTYGTYGTTTTGGGTTGTATTG-3' |
| CTCF1 R1 | 5'-CAACATTCCYGACTACAAAAACCCTT-3' |
| CTCF2 F1 | 5'-TTYGGTTAGGTTGAGGTTT-3' |
| CTCF2 R1 | 5'-TTAACAAAAACAAATTTCCC-3' |
| CTCF1 F Miseq | 5'-TCGTCGGCAGCGTCAGATGTGTATAAGAGACAGGTTGTATTGGGTTGGTGGTTTA-3' |
| CTCF1 R Miseq | 5'- GTCTCGTGGGCTCGGAGATGTGTATAAGAGACAG CTACAAAAACCCTTYGAACCC-3' |
| CTCF2 F Miseq | 5'-TCGTCGGCAGCGTCAGATGTGTATAAGAGACAGTAAATTGTAGGTTTGGGAAG-3' |
| CTCF2 R Miseq | 5'-GTCTCGTGGGCTCGGAGATGTGTATAAGAGACAGTTAACAAAAACAAATTTCCC-3' |

| target name | reference number |
|---|---|

| | |
|---|---|
| mouse anti-MSH2 | Merck Millipore nr NA27 |
| mouse anti-MSH6 | BD transduction Laboratories nr 610919 |
| rabbit anti-MSH3 | Abcam ab154521 |
| mouse anti-Actin ab-5 | BD transduction Laboratories nr 612657 |
| donkey anti-mouse IRDye 800CW | LI-COR nr 925-32212 |
| donkey anti-rabbit IRDye 800CW | LI-COR nr 926-32213 |

**Figure S1.** The *MSH2* exon 3 genomic region edited with CRISPR/Cas9 (Cas) and CRISPR/Cas9Nick (nick). Indicated are the CRISPR/Cas9 and CRISPR/Cas9Nick guide RNAs with their respective PAM sequence. The cutting sites of Cas9 or Cas9Nick are indicated with a red triangle. *MSH2* exon3 F and R indicate the PCR primers used to determine the mutation.

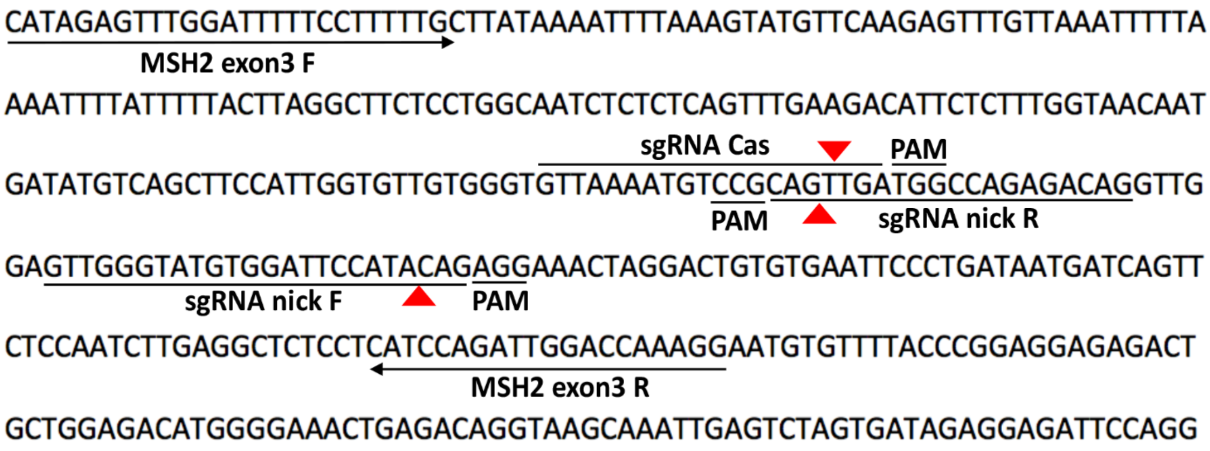


**Figure S2. Mutations and characterization of the clonal MSH2KD lines obtained from VUB03-DM1 and VUB19-DM1 hESCs lines.**

(A) *MSH2* mutations induced at the genome, cDNA and protein level.

(B) *MSH2, MSH3* and *MSH6* mRNA expression analysis for all MSH2WT and MSH2KD DM1 clonal lines. Messenger RNA expression is presented as a fold change per clonal line and relative to one MSH2WT subline set to 1. *UBC*, *GAPDH* and *GUSB* were used as endogenous controls (n=1).

(C) Immunocytochemistry for MSH2 protein in MSH2WT and MSH2KD clonal lines.

A

| cell lines | mutation type | coding effect | genome mutation | cDNA mutation | protein mutation | GT |
|---|---|---|---|---|---|---|
| VUB03KD-SC1 | duplication | stop gain | Chr2(GRCh37):g.47637331dup | NM_000251.2:c.465dup | p.Asp156* | 0/1 |
| | deletion | frameshift | Chr2(GRCh37): g.47637330_47637351del | NM_000251.2:c.464_485del | p.Val155Glufs*12 | 0/1 |
| VUB03KD-SC2 | deletion | in-frame | Chr2(GRCh37):g.47637343_47637378del | NM_000251.2:c.477_512del | p.Gln160_Arg171del | 1/1 |
| VUB03KD-SC3 | deletion | in-frame | Chr2(GRCh37):g.47637343_47637378del | NM_000251.2:c.477_512del | p.Gln160_Arg171del | 1/1 |
| VUB19KD-SC1 | delins | frameshift | Chr2(GRCh37):g.47637366_47637373delins70 | NM_00251.2:c.500_507delins70 | Asp167Valfs*28 | 0/1 |
| | duplication | frameshift | Chr2(GRCh37):g.47637368_47637372dup | NM_000251.2:c.502_506dup | p.Gln170Profs*6 | 0/1 |

B

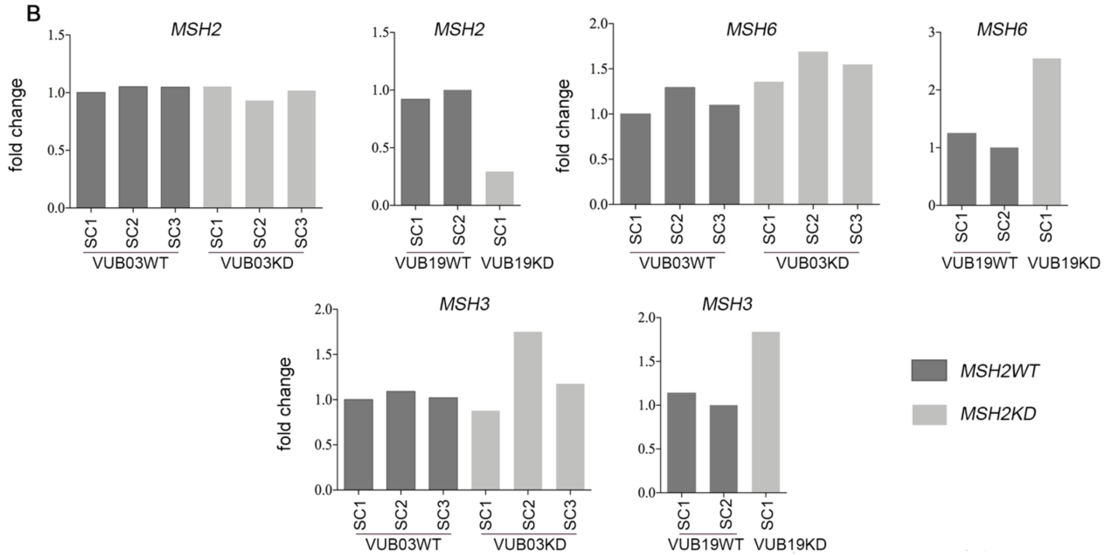


C

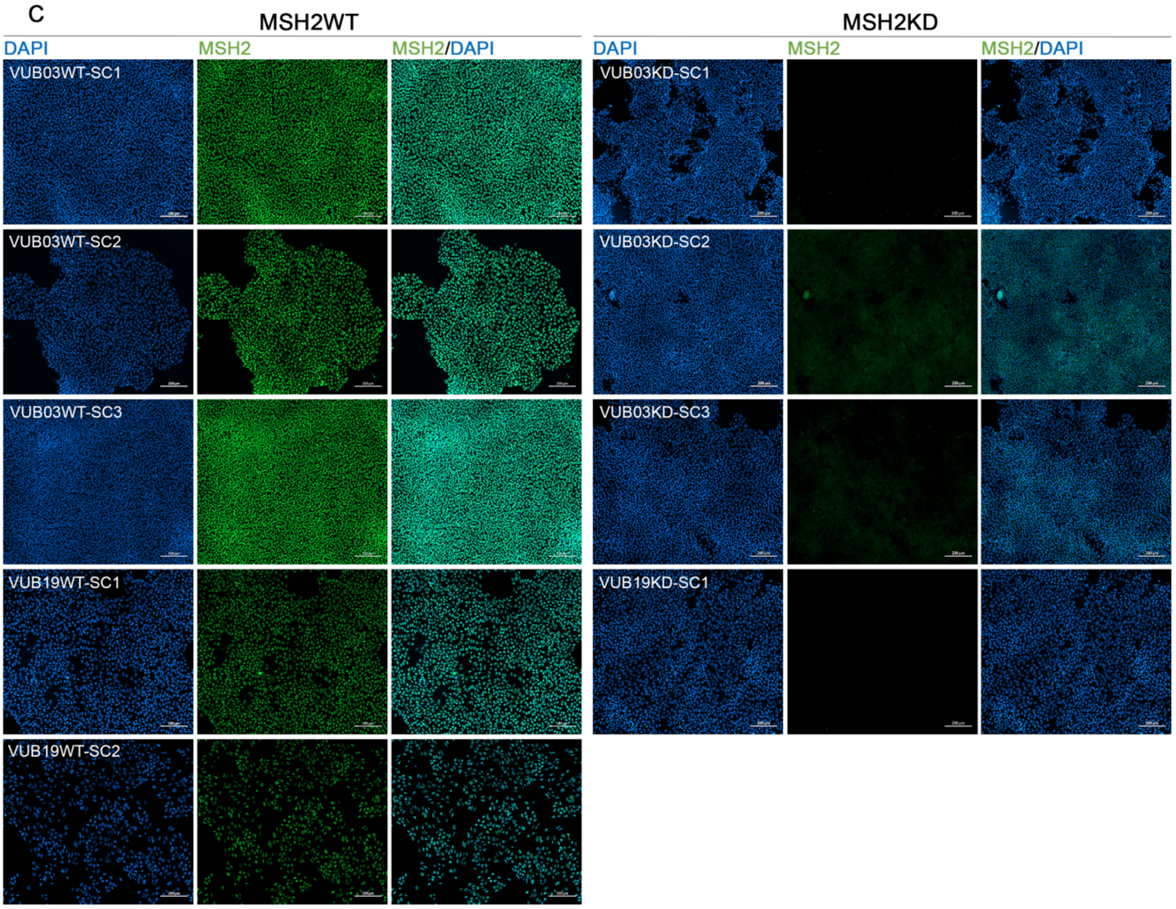

**Figure S3. Mutations and characterization of the clonal MSH2KD lines obtained from non-DM1 VUB02 and VUB06 hESCs lines.**

(A) *MSH2* mutations induced at the genome, cDNA and protein level.

(B) *MSH2, MSH3* and *MSH6* mRNA expression analysis for all MSH2WT and MSH2KD control clonal lines. Messenger RNA expression is presented as a fold change per clonal lines and relative to one MSH2WT subline set to 1. *UBC*, *GAPDH* and *GUSB* were used as endogenous controls (n=1).

(C) Immunocytochemistry for MSH2 protein in MSH2WT and MSH2KD clonal lines.

A

| cell lines | mutation type | coding effect | genome mutation | cDNA mutation | protein mutation | GT |
|---|---|---|---|---|---|---|
| VUB02KD-SC1 | deletion | frameshift | Chr2(GRCh37):g.47637323_47637335del | NM_000251.2:c.457_469del | p.Ser153Alafs*17 | 0/1 |
| | deletion | stop gain | Chr2(GRCh37):g.47637328_47637389del | NM_000251.2:c.462_523del | p.Asp156* | 0/1 |
| VUB02KD-SC2 | delins | frameshift | Chr2(GRCh37): g.47637330_47637368delins74 | NM_000251.2: c.464_502delins74 | p.Val155Glufs*31 | 0/1 |
| | deletion | frameshift | Chr2(GRCh37):g.47637338_47637375del | NM_000251.2:c.472_509del | p.Gln158Glufs*7 | |
| VUB06KD-SC1 | deletion | frameshift | Chr2(GRCh37):g.47637353_47637380del | NM_000251.2:c.487_514del | p.Val163Asnfs*2 | 0/1 |
| | deletion | frameshift | Chr2(GRCh37):g.47637354_47637378del | NM_000251.2:c.488_512del | p.Val163Glyfs*3 | |
| VUB06KD-SC2 | delins | frameshift | Chr2(GRCh37):g.47637343_47637358delinsGT | NM_000152.2: c.477_492delinsGT | p.Gln160Leufs*13 | 0/1 |
| | duplication | frameshift | Chr2(GRCh37):g.47637370_47637374dup | NM_000251.2:c.504_508dup | p.Gln170Profs*6 | |

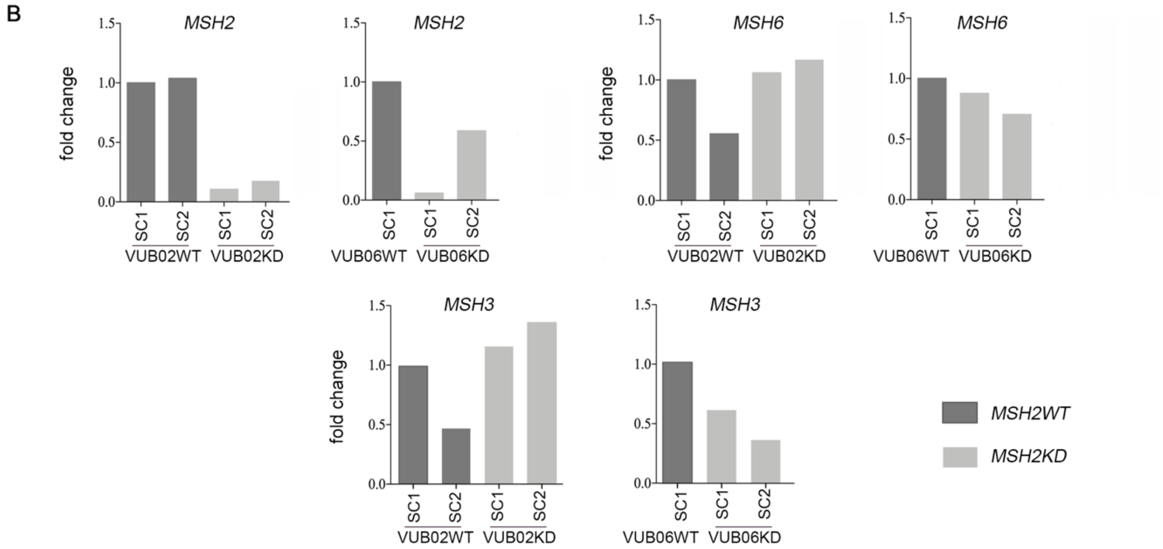


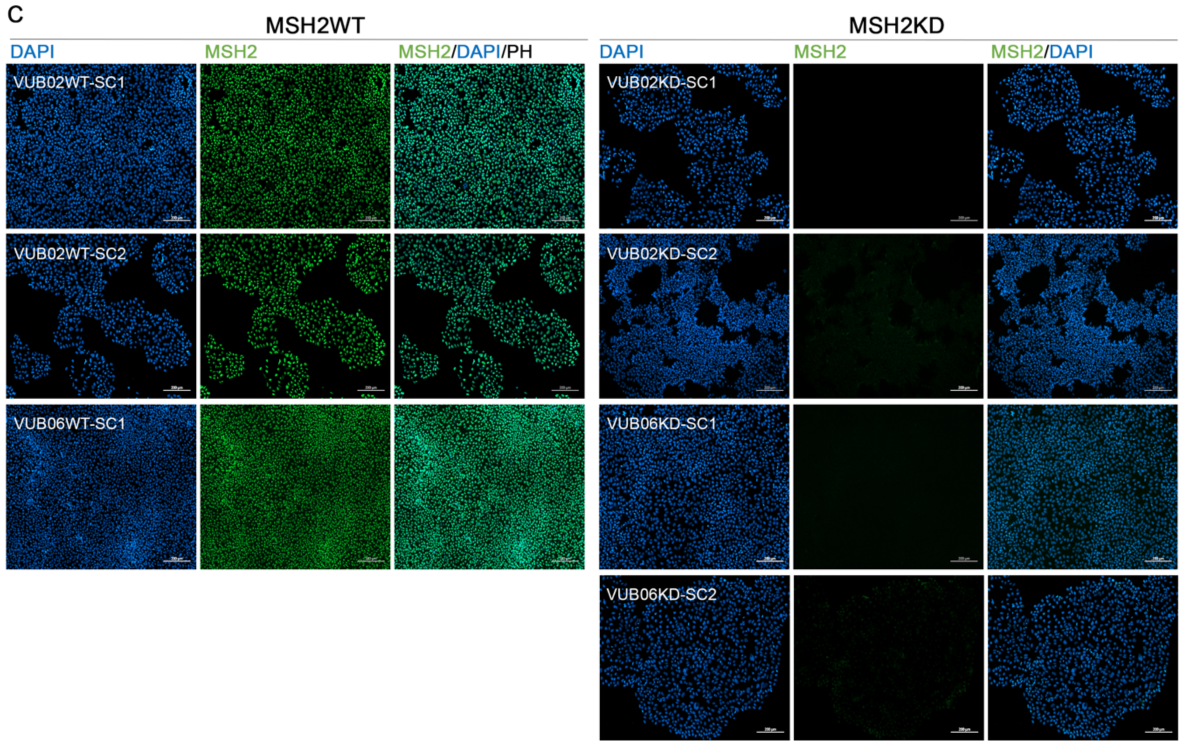

**Figure S4. MSH3 western blot and protein quantifications of MSH2, MSH6 and MSH3.**

(A) Western blot for MSH3 of all DM1 MSH2WT and MSH2KD cell lines. ACTIN was used as endogenous protein loading control.

(B) Western blot for MSH3 of non-DM1 MSH2WT and MSH2KD cell lines. ACTIN was used as endogenous protein loading control.

(C) Protein quantification of MSH2 in DM1 cell lines (VUB03WT, VUB03KD, VUB19WT and VUB19KD).

(D) Protein quantification of MSH2 in non-DM1 cell lines (VUB02WT, VUB02KD, VUB06WT and VUB06KD).

(E) Protein quantification of MSH6 in DM1 cell lines (VUB03WT, VUB03KD, VUB19WT and VUB19KD).

(F) Protein quantification of MSH6 in non-DM1 cell lines (VUB02WT, VUB02KD, VUB06WT and VUB06KD).

(G) Protein quantification of MSH3 in DM1 cell lines (VUB03WT, VUB03KD, VUB19WT and VUB19KD).

(H) Protein quantification of MSH3 in non-DM1 cell lines (VUB02WT, VUB02KD, VUB06WT and VUB06KD).

ACTIN was used as endogenous control and samples are presented in the graph as relative to one MSH2WT subline (n=2, t-test).

Abbreviations: SC: single cell clonal line, WT: *MSH2* wild type, KD: *MSH2* knock-down

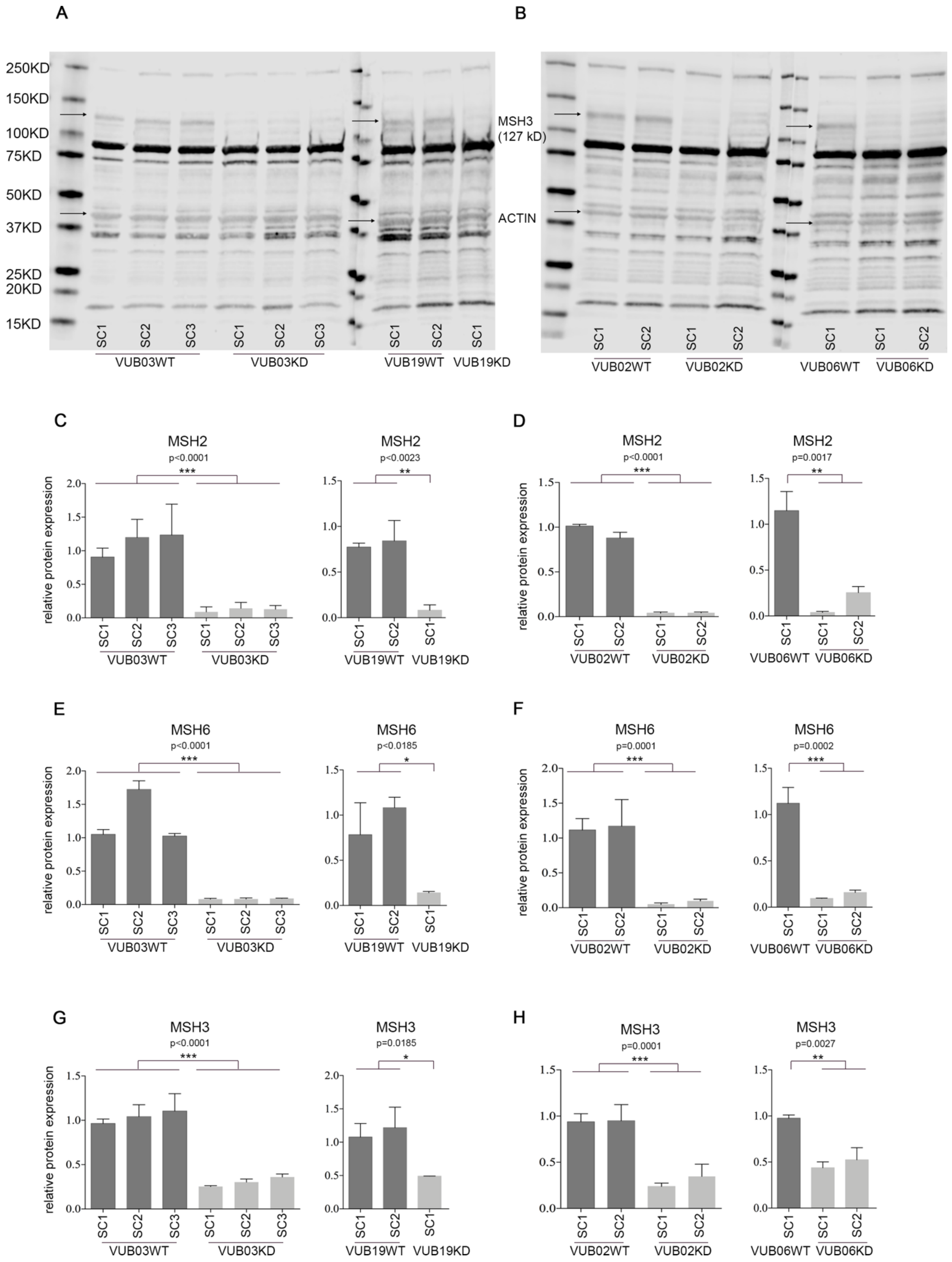
A
B
250KD
150KD
100KD
75KD
50KD
37KD
25KD
20KD
15KD
MSH3
(127 kD)
ACTIN
SC1 SC2 SC3 SC1 SC2 SC3 SC1 SC2 SC1
VUB03WT VUB03KD VUB19WT VUB19KD
SC1 SC2 SC1 SC2 SC1 SC1 SC2
VUB02WT VUB02KD VUB06WT VUB06KD
C
MSH2
p<0.0001
***
MSH2
p<0.0023
**
D
MSH2
p<0.0001
***
MSH2
p=0.0017
**
E
MSH6
p<0.0001
***
MSH6
p<0.0185
*
F
MSH6
p=0.0001
***
MSH6
p=0.0002
***
G
MSH3
p<0.0001
***
MSH3
p=0.0185
*
H
MSH3
p=0.0001
***
MSH3
p=0.0027
**
relative protein expression

**Table S2. The Jonckheere-Terpstra Test details.**

The Jonckheere-Terpstra test was performed on every clonal line, including passages 4, 12 and 20 as 3 different levels. P=passage; N is the number of data points in the sample.

| VUB03WT-SC1 MSH2WT | |
|---|---|
| p4-p12-p20 | |
| | data |
| number of levels | 3 |
| N | 34 |
| Observed J-T Statistic | 289 |
| Mean J-T statistic | 168 |
| std deviation of J-T statistic | 30.121 |
| std J-T statistic | 4.016 |
| Asymp. Sig. (2-tailed) | 0 |

| VUB03WT-SC2 MSH2WT | |
|---|---|
| p4-p12-p20 | |
| | data |
| number of levels | 3 |
| N | 48 |
| Observed J-T Statistic | 728 |
| Mean J-T statistic | 382.5 |
| std deviation of J-T statistic | 52.785 |
| std J-T statistic | 6.545 |
| Asymp. Sig. (2-tailed) | 0 |

| VUB03WT-SC3 MSH2WT | |
|---|---|
| p4-p12-p20 | |
| | data |
| number of levels | 3 |
| N | 22 |
| Observed J-T Statistic | 140 |
| Mean J-T statistic | 78 |
| std deviation of J-T statistic | 16.401 |
| std J-T statistic | 3.78 |
| Asymp. Sig. (2-tailed) | 0 |

| VUB03KD-SC1 MSH2KD | |
|---|---|
| p4-p12-p20 | |
| | data |
| number of levels | 3 |
| N | 36 |
| Observed J-T Statistic | 21 |
| Mean J-T statistic | 214 |
| std deviation of J-T statistic | 34.336 |
| std J-T statistic | -5.621 |
| Asymp. Sig. (2-tailed) | 0 |

| VUB03KD-SC2 MSH2KD | |
|---|---|
| p4-p12-p20 | |
| | data |
| number of levels | 3 |
| N | 51 |
| Observed J-T Statistic | 71,5 |
| Mean J-T statistic | 429 |
| std deviation of J-T statistic | 57.627 |
| std J-T statistic | -6.204 |
| Asymp. Sig. (2-tailed) | 0 |

| VUB03KD-SC3 MSH2KD | |
|---|---|
| p4-p12-p20 | |
| | data |
| number of levels | 3 |
| N | 38 |
| Observed J-T Statistic | 22 |
| Mean J-T statistic | 238,5 |
| std deviation of J-T statistic | 37.212 |
| std J-T statistic | -5.818 |
| Asymp. Sig. (2-tailed) | 0 |

| VUB19WT-SC1 MSH2WT | |
|---|---|
| p4-p12-p20 | |
| | data |
| number of levels | 3 |
| N | 53 |
| Observed J-T Statistic | 905,5 |
| Mean J-T statistic | 465,5 |
| std deviation of J-T statistic | 61.133 |
| std J-T statistic | 7.197 |
| Asymp. Sig. (2-tailed) | 0 |

| VUB19WT-SC2 MSH2WT | |
|---|---|
| p4-p12-p20 | |
| | data |
| number of levels | 3 |
| N | 48 |
| Observed J-T Statistic | 778 |
| Mean J-T statistic | 400 |
| std deviation of J-T statistic | 54.499 |
| std J-T statistic | 6.936 |
| Asymp. Sig. (2-tailed) | 0 |

| VUB19KD-SC1 MSH2KD | |
|---|---|
| p4-p12-p20 | |
| | data |
| number of levels | 3 |
| N | 44 |
| Observed J-T Statistic | 79,5 |
| Mean J-T statistic | 315,5 |
| std deviation of J-T statistic | 45.997 |
| std J-T statistic | -5.131 |
| Asymp. Sig. (2-tailed) | 0 |

**Table S3. Individual CTG repeat values for the right panel of Figures 2 and 3**. PacBio results are shown per individual LongAmp PCR reaction and per single cell clonal line. Number of reads, maximum and minimum repeat size, median and mean CTG repeat sizes are shown per PCR reaction. All values for MSH2WT or MSH2KD clonal lines from VUB03-DM1 and VUB19-DM1 are shown for passages 4, 12 and 20 (**Figure 2**). The values at the bottom of the rows with the mean and median per PCR reaction indicate the mean or median for the whole sample. Values for all *MSH2* transgenic clonal lines for VUB03-DM1 are shown for passage 29. The values at the bottom of the rows with the mean and median per PCR reaction indicate the mean or median respectively for the whole sample (**Figure 3**).

**Table S4. Epi-allele percentages for VUB03-DM1 and VUB19-DM1 at different passages for the MSH2WT and MSH2KD sublines.** A total of 100 epi-alleles were analyzed, covering 25 CpGs upstream of the CTG repeat and 11 CpGs downstream. The methylation percentage upstream (A,B,E,F) and downstream (C,D,G,H) for VUB03-DM1 (p4, p8, p12, p16, p20) and VUB19-DM1 (p4, p8, p12, p16, p20) clonal lines are shown.

101

**A. VUB03-DM1 MSH2WT, upstream**

| | VUB03WT-SC1 | | | | | VUB03WT-SC2 | | | | | VUB03WT-SC3 | | | | |
|---|---|---|---|---|---|---|---|---|---|---|---|---|---|---|---|
| | p4 | p8 | p12 | p16 | p20 | p4 | p8 | p12 | p16 | p20 | p4 | p8 | p12 | p16 | p20 |
| methylation | 84% | 78% | 74% | 66% | 62% | 48% | 69% | 46% | 73% | 74% | 83% | 82% | 63% | 69% | 56% |
| no methylation | 16% | 22% | 26% | 34% | 38% | 52% | 31% | 54% | 27% | 26% | 17% | 18% | 37% | 31% | 44% |

**B. VUB03-DM1 MSH2KD, upstream**

| | VUB03KD-SC1 | | | | | VUB03KD-SC2 | | | | | VUB03KD-SC3 | | | | |
|---|---|---|---|---|---|---|---|---|---|---|---|---|---|---|---|
| | p4 | p8 | p12 | p16 | p20 | p4 | p8 | p12 | p16 | p20 | p4 | p8 | p12 | p16 | p20 |
| methylation | 39% | 11% | 17% | 17% | 8% | 12% | 15% | 19% | 20% | 30% | 19% | 12% | 10% | 9% | 0% |
| no methylation | 61% | 89% | 83% | 83% | 92% | 88% | 85% | 81% | 80% | 70% | 81% | 88% | 90% | 91% | 100% |

**C. VUB03-DM1 MSH2WT, downstream**

| | VUB03WT-SC1 | | | | | VUB03WT-SC2 | | | | | VUB03WT-SC3 | | | | |
|---|---|---|---|---|---|---|---|---|---|---|---|---|---|---|---|
| | p4 | p8 | p12 | p16 | p20 | p4 | p8 | p12 | p16 | p20 | p4 | p8 | p12 | p16 | p20 |
| methylation | 72% | 84% | 85% | 90% | 90% | 79% | 93% | 76% | 85% | 90% | 67% | 86% | 76% | 86% | 91% |
| no methylation | 28% | 16% | 15% | 10% | 10% | 21% | 7% | 24% | 15% | 10% | 33% | 14% | 24% | 14% | 9% |

**D. VUB03-DM1 MSH2KD, downstream**

| | VUB03KD-SC1 | | | | | VUB03KD-SC2 | | | | | VUB03KD-SC3 | | | | |
|---|---|---|---|---|---|---|---|---|---|---|---|---|---|---|---|
| | p4 | p8 | p12 | p16 | p20 | p4 | p8 | p12 | p16 | p20 | p4 | p8 | p12 | p16 | p20 |
| methylation | 66% | 83% | 88% | 82% | 85% | 73% | 76% | 84% | 83% | 81% | 72% | 83% | 88% | 89% | 88% |
| no methylation | 34% | 16% | 12% | 18% | 15% | 27% | 24% | 16% | 17% | 19% | 28% | 17% | 13% | 11% | 12% |

**E. VUB19-DM1 MSH2WT, upstream**

| | VUB19WT-SC1 | | | | | VUB19WT-SC2 | | | | |
|---|---|---|---|---|---|---|---|---|---|---|
| | p4 | p8 | p12 | p16 | p20 | p4 | p8 | p12 | p16 | p20 |
| methylation | 9% | 0% | 18% | 9% | 26% | 25% | 28% | 14% | 33% | 56% |
| no methylation | 91% | 100% | 82% | 91% | 74% | 75% | 72% | 86% | 67% | 44% |

**F. VUB19-DM1 MSH2KD, upstream**

| | VUB19KD-SC1 | | | | |
|---|---|---|---|---|---|
| | p4 | p8 | p12 | p16 | p20 |
| methylation | 0% | 0% | 0% | 0% | 1% |
| no methylation | 100% | 100% | 100% | 100% | 99% |

**G. VUB19-DM1 MSH2WT, downstream**

| | VUB19WT-SC1 | | | | | VUB19WT-SC2 | | | | |
|---|---|---|---|---|---|---|---|---|---|---|
| | p4 | p8 | p12 | p16 | p20 | p4 | p8 | p12 | p16 | p20 |
| methylation | 87% | 87% | 89% | 80% | 71% | 85% | 91% | 74% | 83% | 77% |
| no methylation | 13% | 13% | 11% | 20% | 29% | 15% | 9% | 26% | 17% | 23% |

**H. VUB19-DM1 MSH2KD, downstream**

| | VUB19KD-SC1 | | | | |
|---|---|---|---|---|---|
| | p4 | p8 | p12 | p16 | p20 |
| methylation | 80% | 85% | 81% | 88% | 81% |
| no methylation | 20% | 15% | 19% | 12% | 19% |

102

**Table S5. Kolmogorov Smirnov test output on the CpG methylation data upstream of the repeat.** MSH2WT and MSH2KD samples were compared at passages 4, 12 and 20 as well as over culture within the same cell lines from p4 to p20. In addition, MSH2TG cell lines at passage 29 were compared to their respective MSH2KD cell lines at passage 12.

| **VUB03WT - VUB03KD** | **VUB03WT - VUB03KD** | **VUB03WT - VUB03KD** |
|---|---|---|
| **p4 WT - p4 KD (pooled)** | **p12 WT - p12 KD (pooled)** | **p20 WT - p20 KD (pooled)** |
| **p=0,025** | **p=0,000** | **p=0,000** |
| data distribution is not the same | data distribution is not the same | data distribution is not the same |

| **VUB03WT** | **VUB03KD** |
|---|---|
| **p4 WT- p20 WT (pooled)** | **p4 KD - p20 KD (pooled)** |
| **p=0,646** | **p=0,448** |
| data distribution is the same | data distribution is the same |

| **VUB19WT - VUB19KD** | **VUB19WT - VUB19KD** | **VUB19WT - VUB19KD** |
|---|---|---|
| **p4 WT - p4 KD (pooled)** | **p12 WT - p12 KD (pooled)** | **p20 WT - p20 KD (pooled)** |
| **p=0,646** | **p=0,172** | **p=0,005** |
| data distribution is the same | data distribution is the same | data distribution is not the same |

| **VUB19WT** | **VUB19KD** |
|---|---|
| **p4 WT- p20 WT (pooled)** | **p4 KD - p20 KD** |
| **p=0,021** | **p=1** |
| data distribution is not the same | data distribution is the same |

| **VUB03TG** | **VUB03TG** | **VUB03TG** | **VUB03TG** |
|---|---|---|---|
| **p12 SC1 KD - p29 SC1 TG** | **p12 SC2 KD - p29 SC2A TG** | **p12 SC2 KD - p29 SC2B TG** | **p12 SC3 KD - p29 SC3 TG** |
| **p=0,206** | **p=0,023** | **p=0,206** | **p=0,206** |
| data distribution is the same | data distribution is not the same | data distribution is the same | data distribution is the same |

**Figure S5. Methylation pattern of the CpG rich region located downstream of the CTG repeat on the *DMPK* locus upon *MSH2* knock-down.** The left panel shows the median CTG repeat size for passage 4, 12 and 20 for clonal lines of VUB03-DM1 and VUB19-DM1 for MSH2WT and MSH2KD. Details see legend Figure 2. The right panel represents the downstream methylation from passage 4 to passage 12 for clonal lines of VUB03-DM1 and VUB19-DM1 for MSH2WT and MSH2KD. The downstream methylation is shown for 11 CpG sites and a total of 100 epi-alleles were analyzed per passage. Details see legend Figure 2. Details of the percentages of methylated or unmethylated epi-alleles are presented in **Table S4**. Abbreviations: pas.: passage, KD: *MSH2* knock-down, WT: *MSH2* wild type

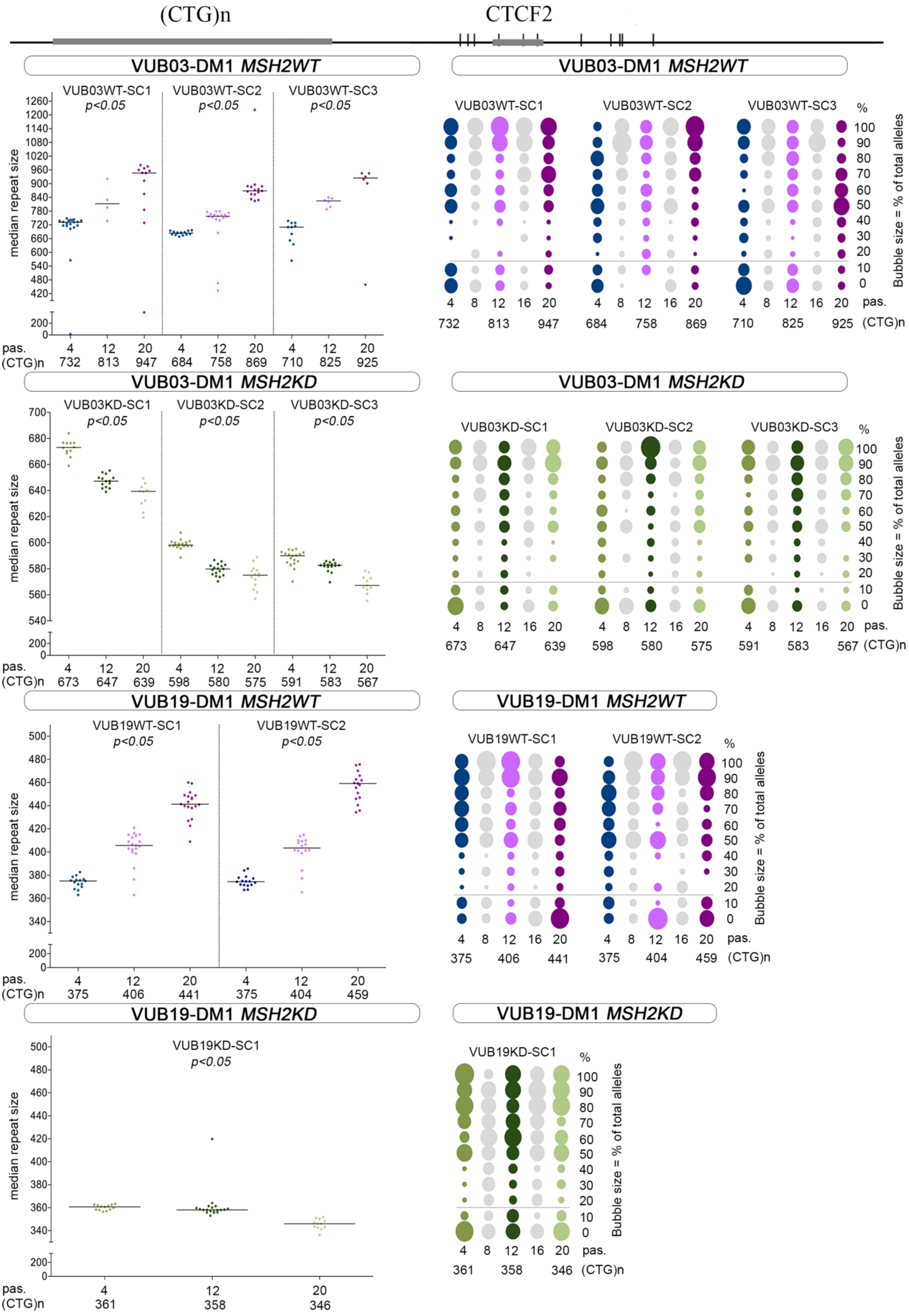

(CTG)n
CTCF2
VUB03-DM1 MSH2WT
VUB03WT-SC1
p<0.05
VUB03WT-SC2
p<0.05
VUB03WT-SC3
p<0.05
median repeat size
pas.
(CTG)n
4 732
12 813
20 947
4 684
12 758
20 869
4 710
12 825
20 925
VUB03-DM1 MSH2KD
VUB03KD-SC1
p<0.05
VUB03KD-SC2
p<0.05
VUB03KD-SC3
p<0.05
4 673
12 647
20 639
4 598
12 580
20 575
4 591
12 583
20 567
VUB19-DM1 MSH2WT
VUB19WT-SC1
p<0.05
VUB19WT-SC2
p<0.05
4 375
12 406
20 441
4 375
12 404
20 459
VUB19-DM1 MSH2KD
VUB19KD-SC1
p<0.05
4 361
12 358
20 346
%
Bubble size = % of total alleles

**Figure S6. Sanger sequences following bisulfite treatment of the CpG site upstream of the repeat of non-DM1 clonal lines VUB02 and VUB06 showing absence of methylation.** CpG sites are indicated in yellow and the converted C to a T is indicated with an arrow.

VUB02WT-SC1 p4 upstream

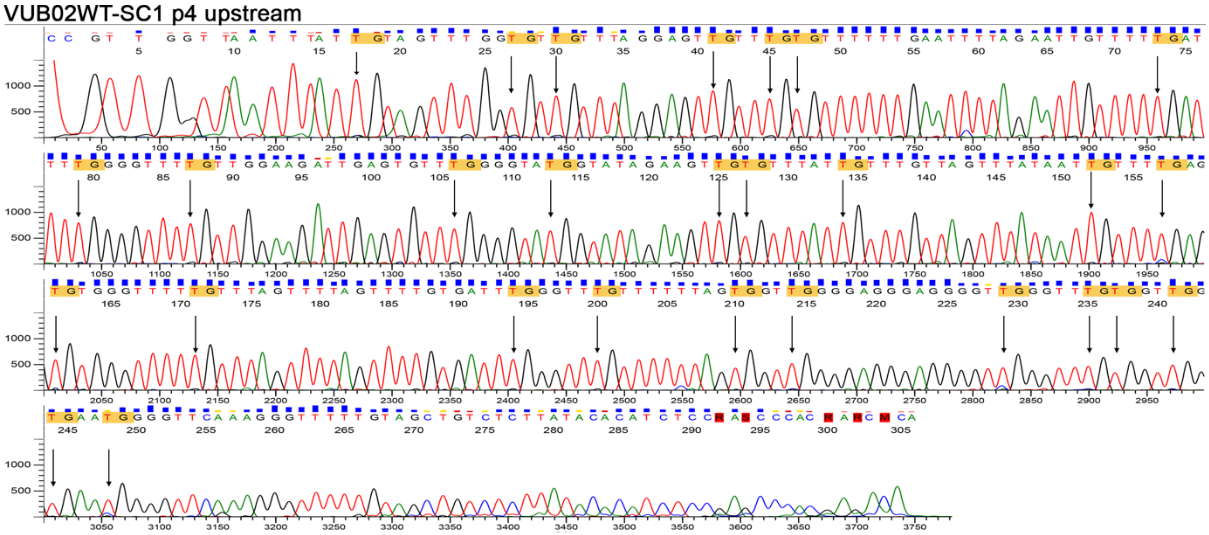

VUB02WT-SC2 p4 upstream

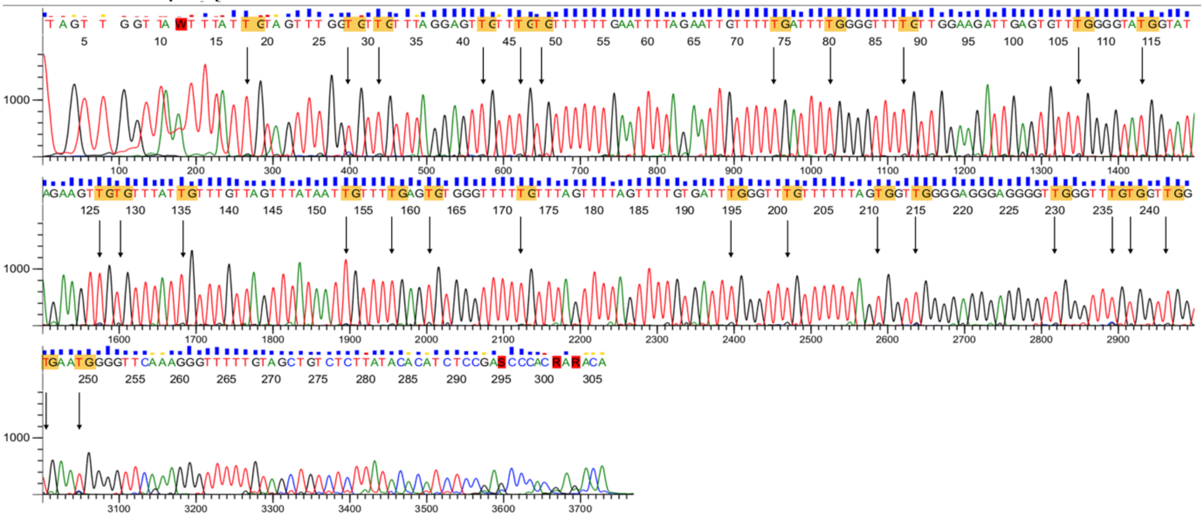

VUB02KD-SC1 p4 upstream

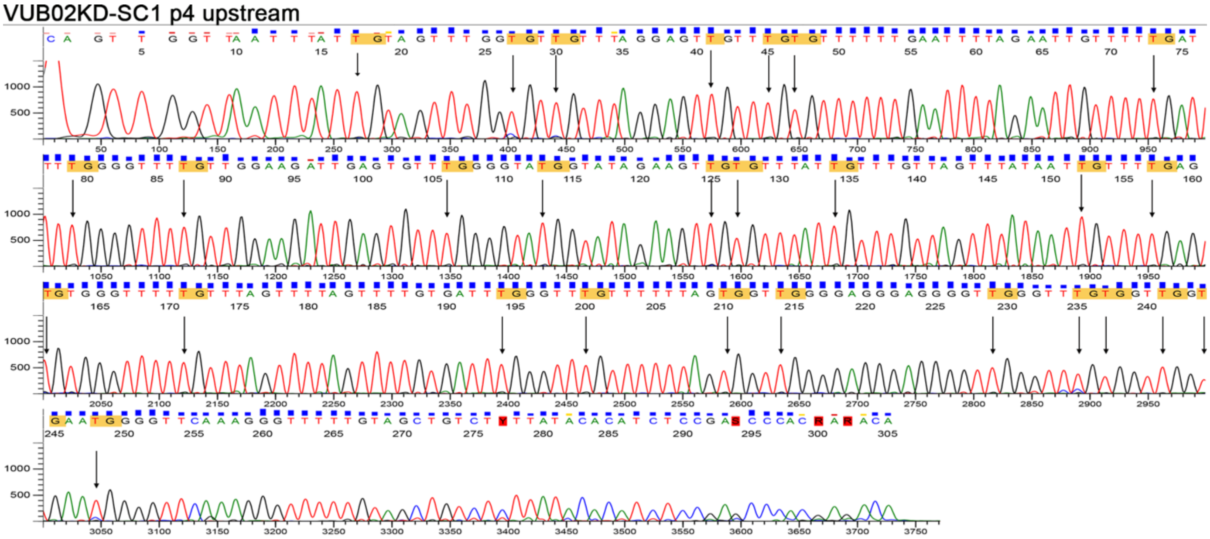

VUB02KD-SC2 p4 upstream

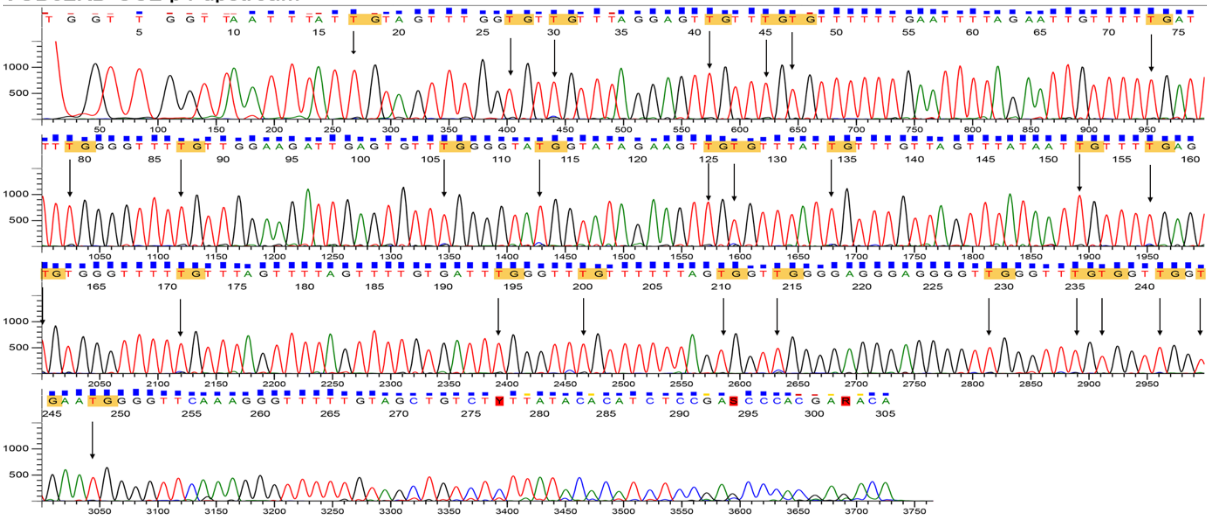

VUB06WT-SC1 p4 upstream

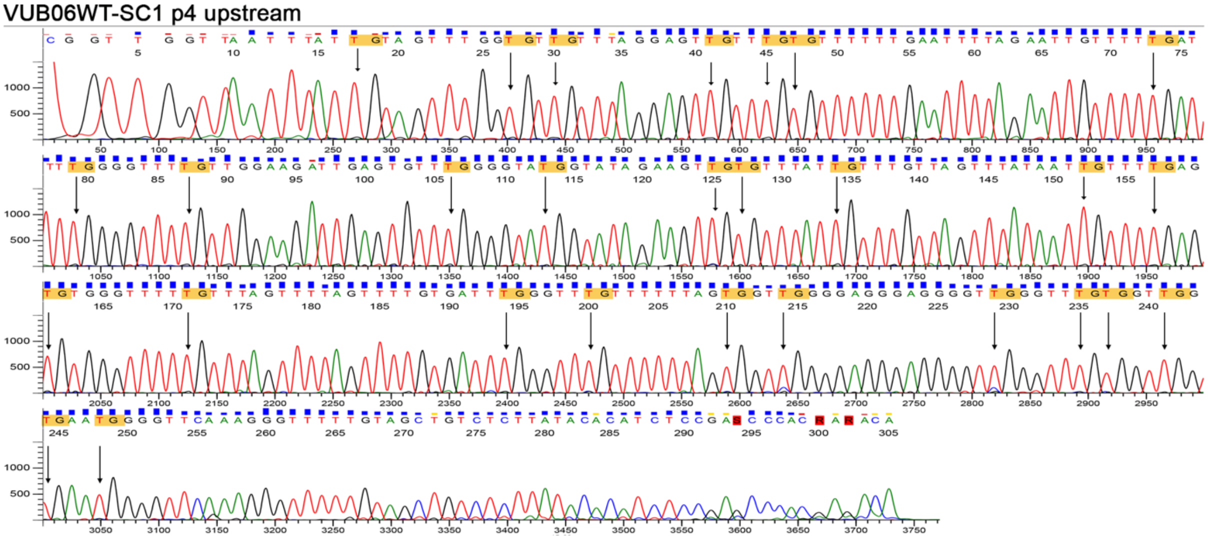

VUB06KD-SC1 p4 upstream

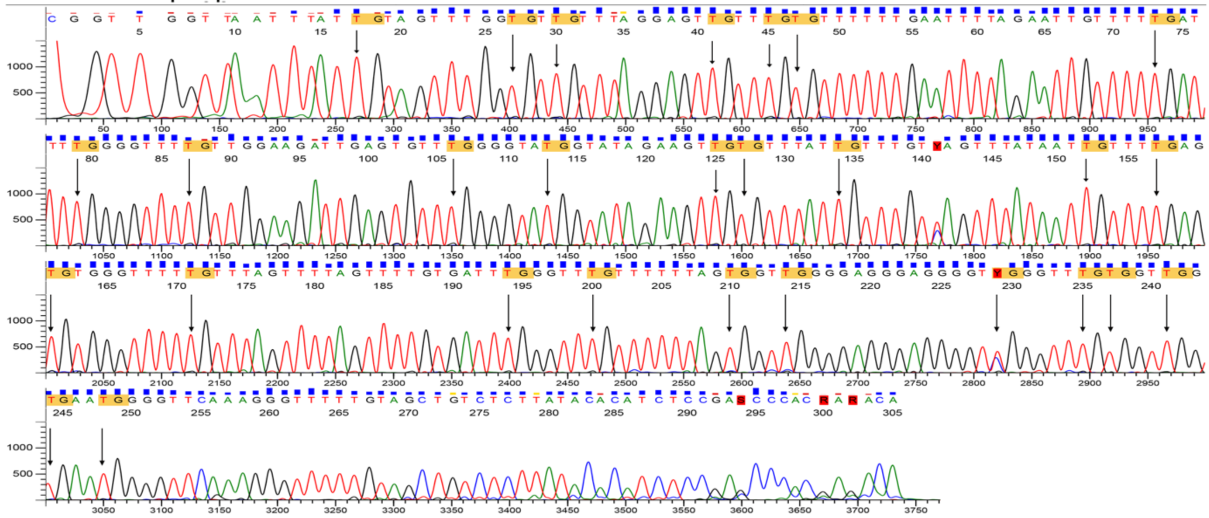

VUB06KD-SC2 p4 upstream

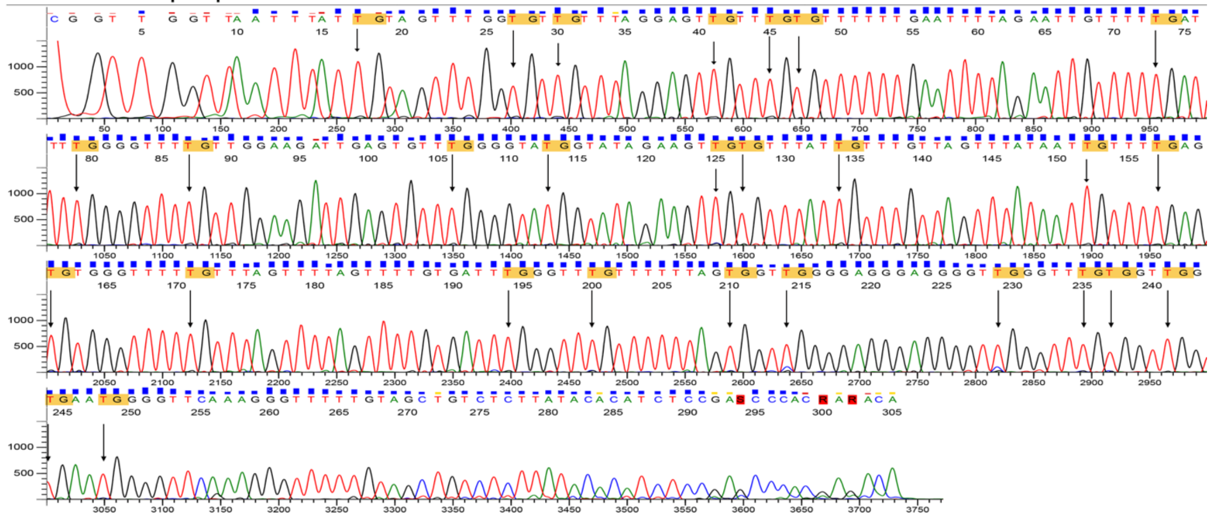

**Figure S7. Sanger sequences following bisulfite treatment of the CpG site downstream of the repeat of non-DM1 clonal lines VUB02 and VUB06 showing absence of methylation.** CpG sites are indicated in yellow and the converted C to a T is indicated with an arrow.

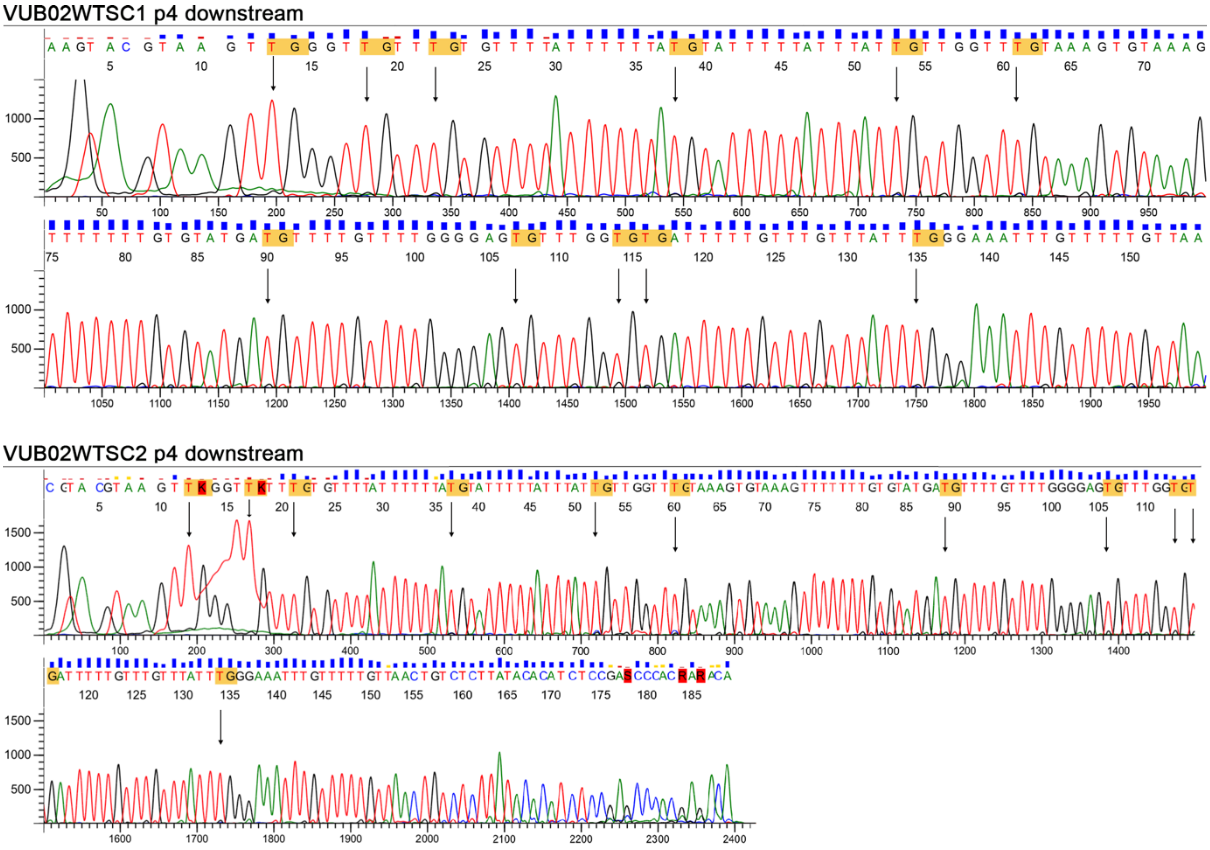

VUB02KD-SC1 p4 downstream

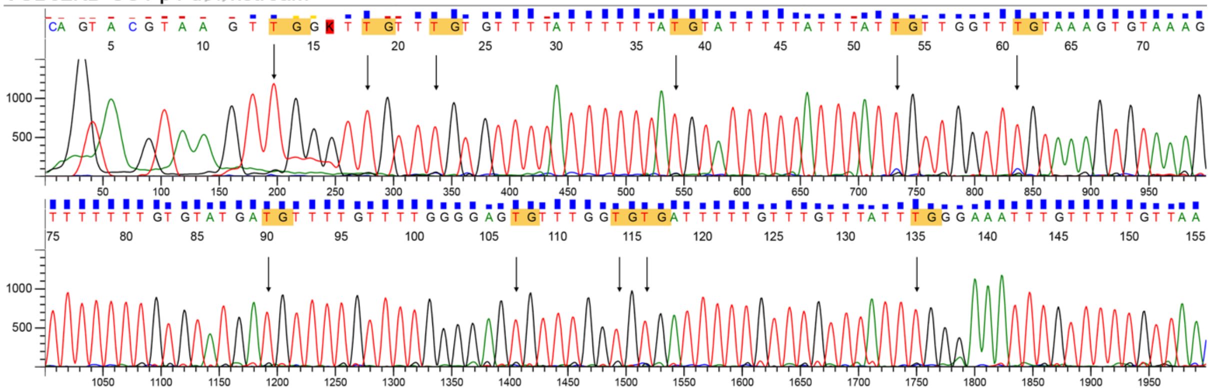

VUB02KD-SC2 p4 downstream

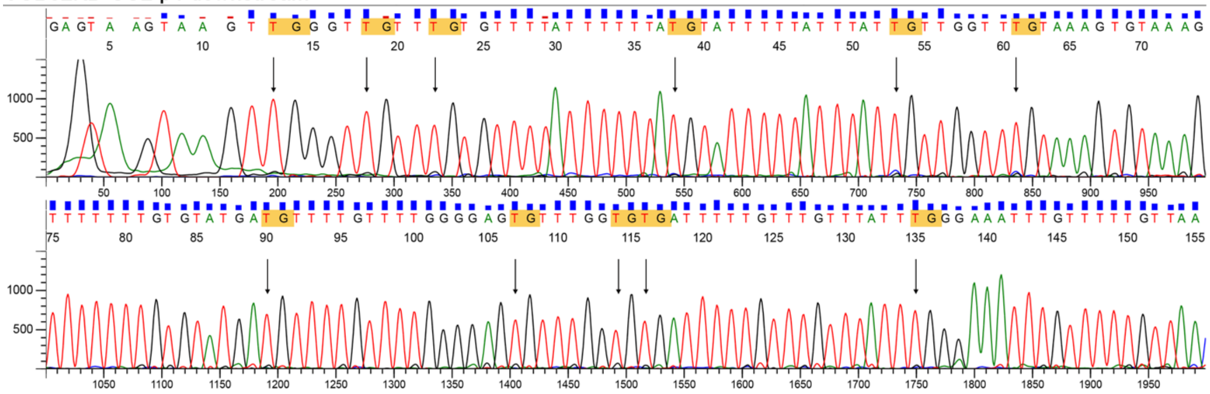

VUB06WT-SC1 p4 downstream

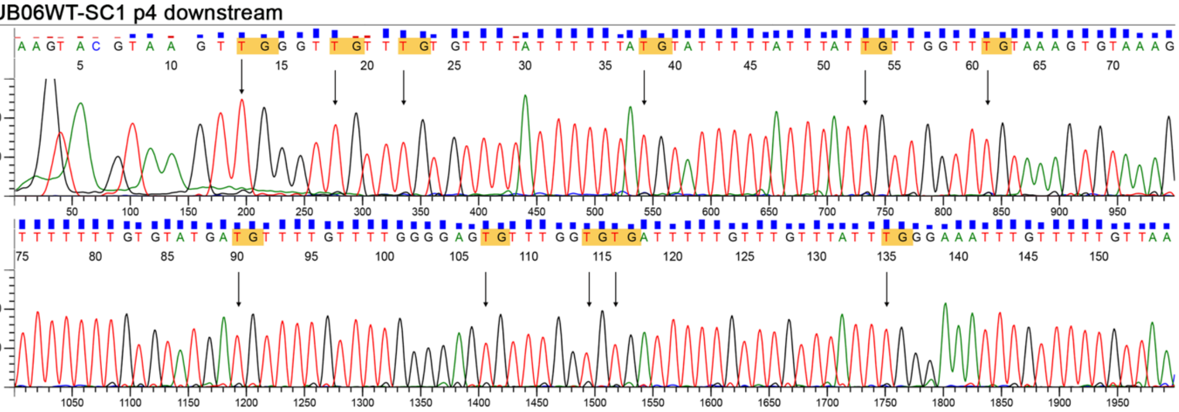

VUB06KD-SC1 p4 downstream

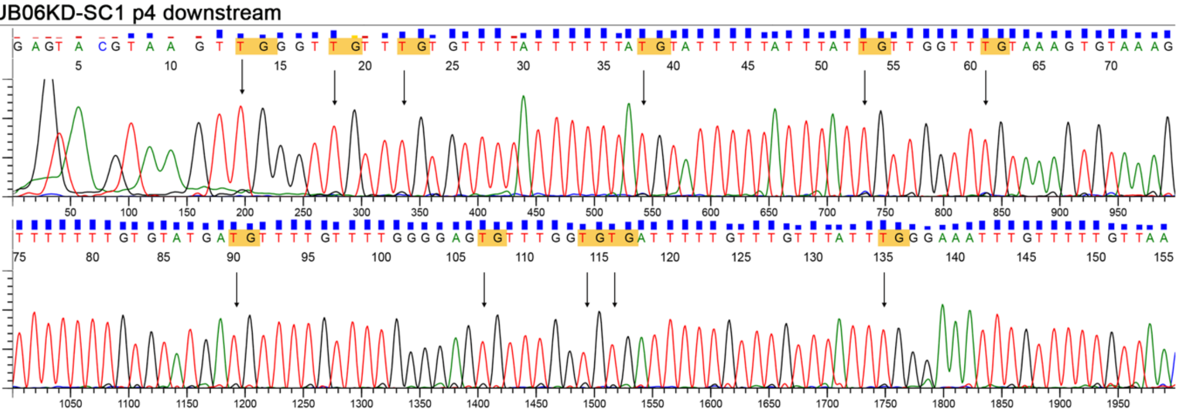

VUB06KD-SC2 p4 downstream

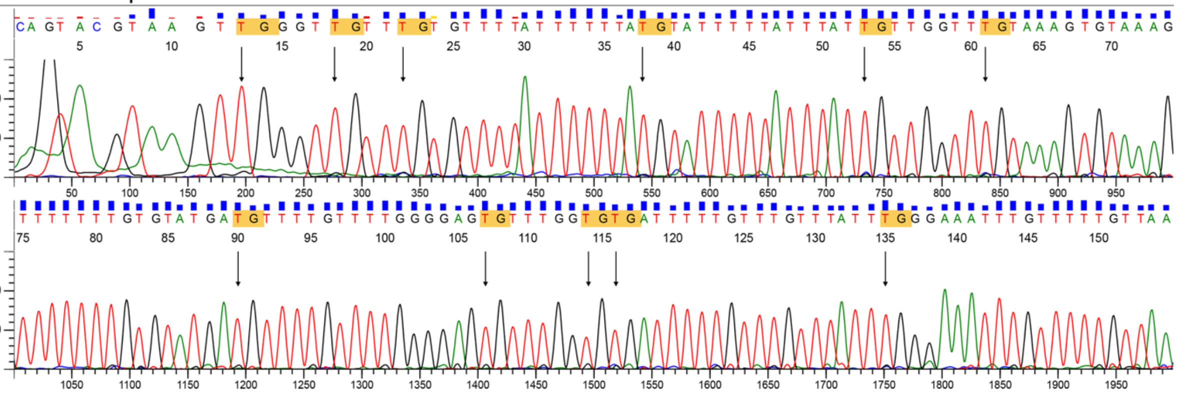

**Figure S8. Immunocytochemistry and Western blot after *MSH2* transgene expression in MSH2KD clonal lines of VUB03-DM1.**

(A) Immunocytochemistry of *MSH2* transgene expression in MSH2KD clonal lines of VUB03-DM1. The MSH2 protein is re-expressed in VUB03TG-SC1, VUB03TG-SC2A, VUB03TG-SC2B, VUB03TG-SC3.

(B) Western blot of MSH2TG lines at passage 29 showing MSH2 and MSH6 proteins (top) and MSH2 only (bottom). ACTIN was used as endogenous control.

(C) *MSH2, MSH3* and *MSH6* mRNA expression analysis for all MSH2WT, MSH2KD and MSH2TG clonal lines of VUB03-DM1. Messenger RNA expression is presented as a fold change per clonal line and relative to one MSH2WT subline set to 1. *UBC*, *GAPDH* and *GUSB* were used as endogenous controls (n=1).

(D) Quantification of MSH2, MSH6 and MSH3 proteins on a Western blot at passage 8 for VUB03WT and VUB03KD cell lines and at passage 29 for VUB03TG cell lines. ACTIN was used as endogenous control and samples are presented in the graph as relative to one MSH2WT subline (n=2, t-test).

Abbreviations: WT: *MSH2* wild type, KD: *MSH2* knock-down, TG: *MSH2* transgene expression, NS: not significant

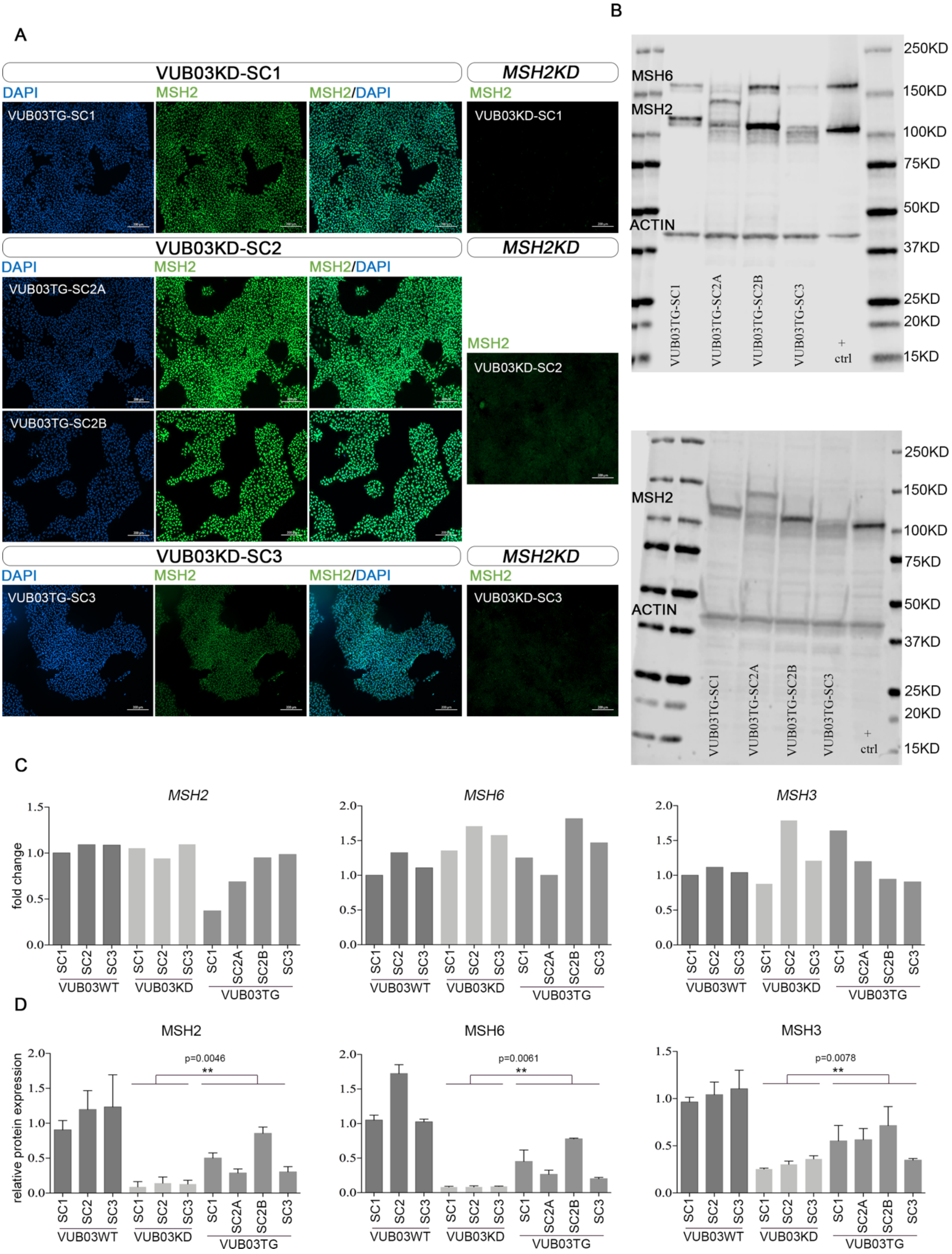
A
VUB03KD-SC1
MSH2KD
DAPI
MSH2
MSH2/DAPI
VUB03TG-SC1
VUB03KD-SC1
VUB03KD-SC2
VUB03TG-SC2A
VUB03TG-SC2B
VUB03KD-SC3
VUB03TG-SC3
B
MSH6
MSH2
ACTIN
250KD
150KD
100KD
75KD
50KD
37KD
25KD
20KD
15KD
VUB03TG-SC1
VUB03TG-SC2A
VUB03TG-SC2B
VUB03TG-SC3
+ ctrl
C
MSH2
MSH6
MSH3
fold change
SC1
SC2
SC3
SC2A
SC2B
VUB03WT
VUB03KD
VUB03TG
D
relative protein expression
p=0.0046
p=0.0061
p=0.0078
**

**Figure S9. Western blot of all VUB03-DM1 lines used in the manuscript, including MSH2WT, MSH2KD and MSH2TG.**

(A) Western blot for MSH2 and MSH6 of all VUB03-DM1 lines used in the manuscript, including MSH2WT at passage 8, MSH2KD at passage 8 and MSH2TG at passage 29. All VUB03-DM1 samples were run on one Western blot gel. The Western blot membrane was divided in two parts to fit a falcon tube and were subsequently re-assembled to scan the Western blot. ACTIN was used as endogenous protein loading control.

(B) Western blot for MSH2 and MSH6 of all VUB03-DM1 MSH2TG lines at passages 21, 25 and 29.

All VUB03-DM1 TG samples at different passages were run on one Western blot gel. The Western blot membrane was divided in two parts to fit a falcon tube and were subsequently re-assembled to scan the Western blot. ACTIN was used as endogenous protein loading control.

(C) Western blot showing the MSH3 protein levels in all MSH2TG. ACTIN was used as endogenous protein loading control.

Abbreviations: p: passage; WT: *MSH2* wild type, KD: *MSH2* knock-down, TG: *MSH2* transgene expression

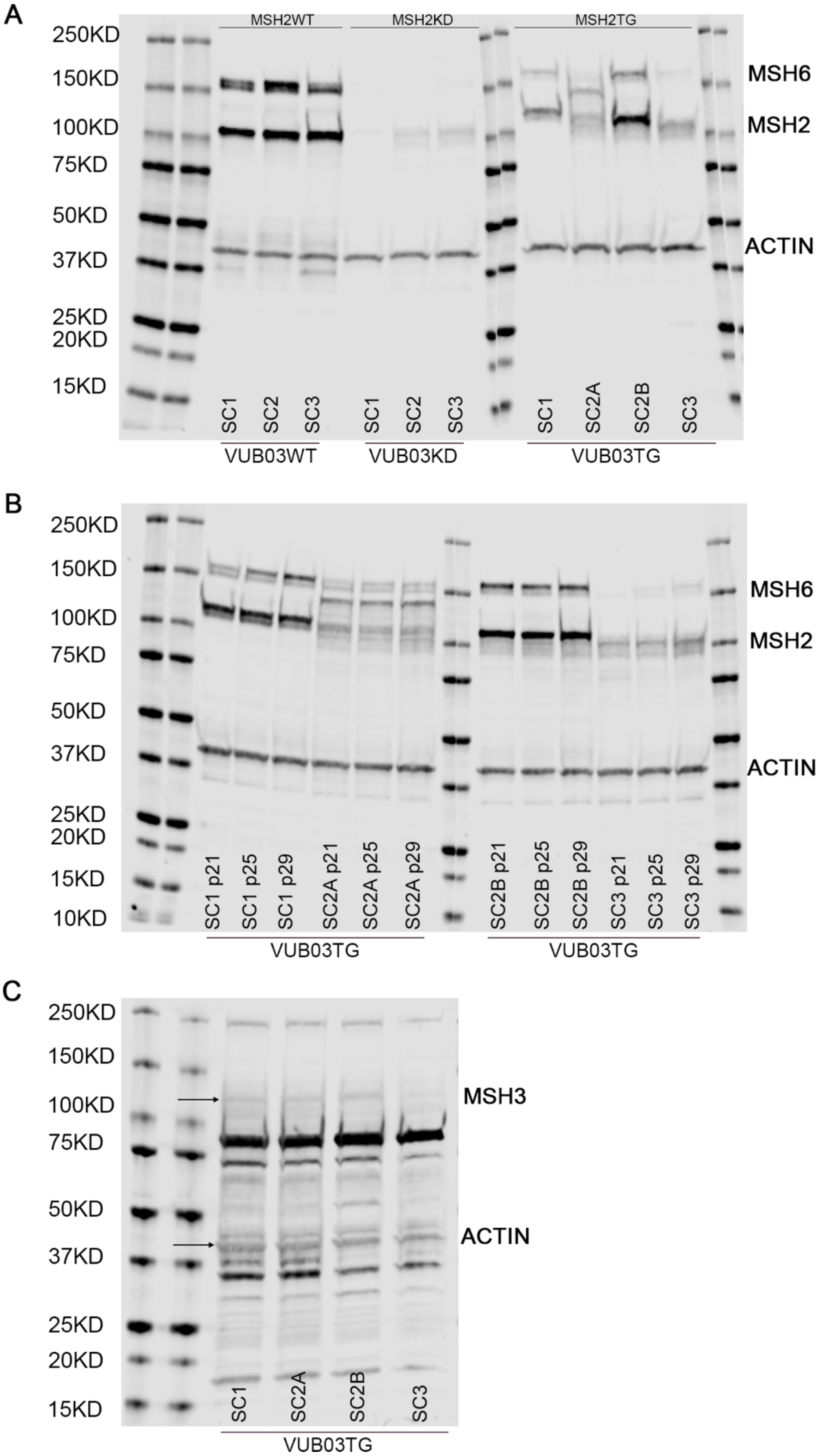
A
250KD
150KD
100KD
75KD
50KD
37KD
25KD
20KD
15KD
MSH2WT
MSH2KD
MSH2TG
MSH6
MSH2
ACTIN
SC1
SC2
SC3
SC1
SC2
SC3
SC1
SC2A
SC2B
SC3
VUB03WT
VUB03KD
VUB03TG
B
250KD
150KD
100KD
75KD
50KD
37KD
25KD
20KD
15KD
10KD
MSH6
MSH2
ACTIN
SC1 p21
SC1 p25
SC1 p29
SC2A p21
SC2A p25
SC2A p29
SC2B p21
SC2B p25
SC2B p29
SC3 p21
SC3 p25
SC3 p29
VUB03TG
VUB03TG
C
250KD
150KD
100KD
75KD
50KD
37KD
25KD
20KD
15KD
MSH3
ACTIN
SC1
SC2A
SC2B
SC3
VUB03TG

**Table S6. Epi-allele percentages before *MSH2* transgene expression, before clonal expansion and after *MSH2* transgene clonal expansion.** A total of 100 epi-alleles were analyzed, covering 25 CpG regions upstream of the CTG repeat and 11 CpG regions downstream. The methylation percentage upstream (A,C,E) and downstream (B,D,F) for VUB03-DM1 (p21, p25, p29) clonal lines are plotted.

**A. VUB03-DM1, upstream**

| | VUB03KD-SC1 | VUB03KD-SC1_TG | VUB03TG-SC1 | | |
|---|---|---|---|---|---|
| | p12 | bulk p19 | p21 | p25 | p29 |
| methylation | 17% | 0% | 37% | 33% | 2% |
| no methylation | 83% | 100% | 63% | 67% | 98% |

**B. VUB03-DM1, downstream**

| | VUB03KD-SC1 | VUB03KD-SC1_TG | VUB03TG-SC1 | | |
|---|---|---|---|---|---|
| | p12 | bulk p19 | p21 | p25 | p29 |
| methylation | 88% | 96% | 81% | 90% | 90% |
| no methylation | 12% | 4% | 19% | 10% | 10% |

**C. VUB03-DM1, upstream**

| | VUB03KD-SC2 | VUB03KD-SC2_TG | VUB03TG-SC2A | | | VUB03TG-SC2B | | |
|---|---|---|---|---|---|---|---|---|
| | p12 | bulk p19 | p21 | p25 | p29 | p21 | p25 | p29 |
| methylation | 19% | 76% | 94% | 71% | 79% | 0% | 0% | 1% |
| no methylation | 81% | 24% | 6% | 29% | 21% | 100% | 100% | 99% |

**D. VUB03-DM1, downstream**

| | VUB03KD-SC2 | VUB03KD-SC2_TG | VUB03TG-SC2A | | | VUB03TG-SC2B | | |
|---|---|---|---|---|---|---|---|---|
| | p12 | bulk p19 | p21 | p25 | p29 | p21 | p25 | p29 |
| methylation | 84% | 90% | 76% | 81% | 89% | 92% | 85% | 90% |
| no methylation | 16% | 10% | 24% | 19% | 11% | 8% | 15% | 10% |

**E. VUB03-DM1, upstream**

| | VUB03KD-SC3 | VUB03KD-SC3_TG | VUB03TG-SC3 | | |
|---|---|---|---|---|---|
| | p12 | bulk p19 | p21 | p25 | p29 |
| methylation | 10% | 0% | 0% | 0% | 2% |
| no methylation | 90% | 100% | 100% | 100% | 98% |

**F. VUB03-DM1, downstream**

| VUB03KD-SC3 | VUB03KD-SC3_TG | VUB03TG-SC3 | | |
|---|---|---|---|---|
| p12 | bulk p19 | p21 | p25 | p29 |

| | | | | | |
|---|---|---|---|---|---|
| methylation | 88% | 85% | 72% | 62% | 95% |
| no methylation | 13% | 15% | 28% | 38% | 5% |

**Figure S10. The methylation pattern of the CpG rich region located downstream of the CTG repeat on the *DMPK* locus after *MSH2* transgene expression.** The left panel shows the median CTG repeat size for passage 12 and 29 for clonal lines of VUB03-DM1. The right panel represents the downstream methylation at passage 12 of MSH2KD and the starting point of *MSH2* transgene expression for clonal lines VUB03KD-SC1, VUB03KD-SC2 and VUB03KD-SC3, as well as passage 21, 25 and 29 after transgene expression for clonal lines VUB03TG-SC1, VUB03TG-SC2A, VUB03TG-SC2B and VUB03TG-SC3. The downstream methylation is shown for 11 CpG sites and a total of 100 epi-alleles were analyzed per passage. The bubble size indicates the percentage of epi-alleles with that specific amount of CpG methylation. The bubbles under the horizontal grey line represent non-methylated alleles and is equivalent to wild type alleles as seen in non-DM1 individuals (Barbé *et al.*, 2017), all bubbles above that line are considered as methylated epi-alleles. CpG sites in the downstream region remain unchanged after *MSH2* re-introduction.

Abbreviations: KD: *MSH2* knock-down, TG: *MSH2* transgene expression

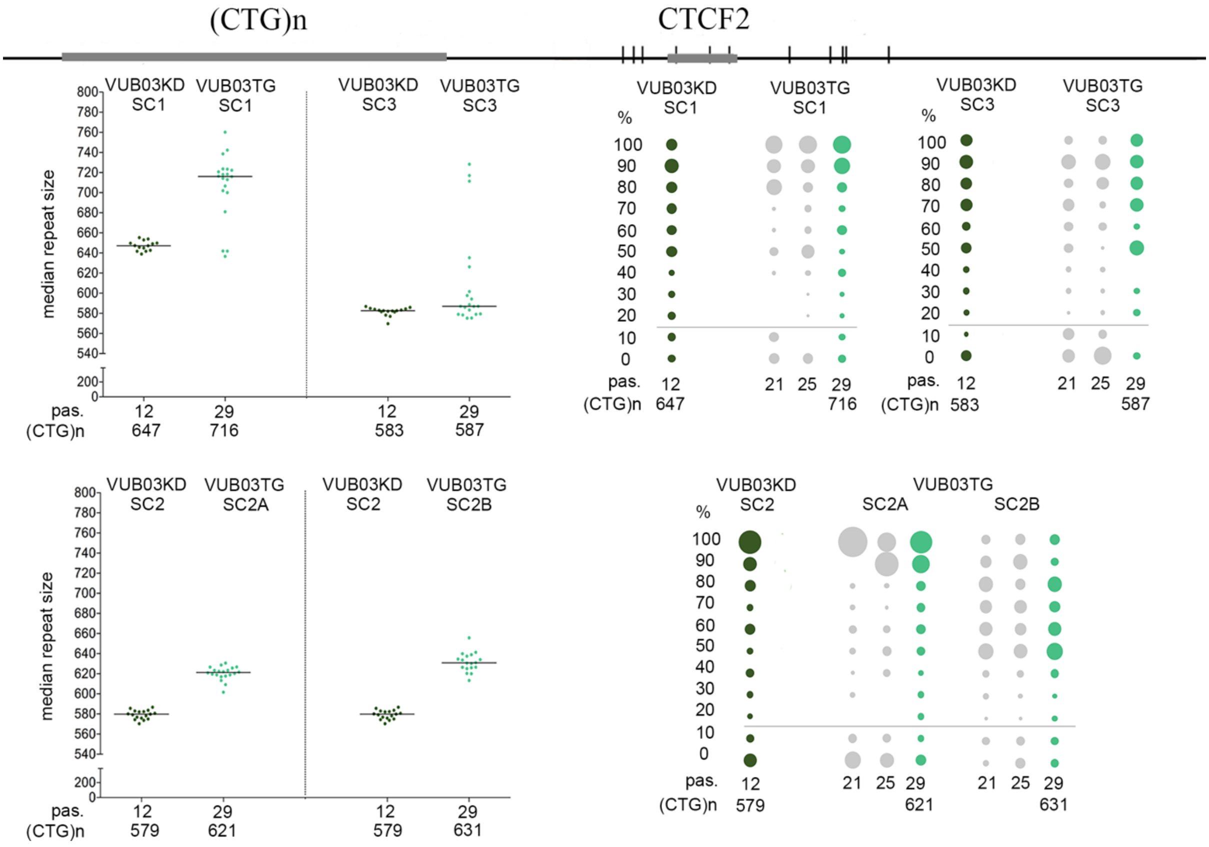
(CTG)n
CTCF2
VUB03KD SC1
VUB03TG SC1
VUB03KD SC3
VUB03TG SC3
median repeat size
pas.
(CTG)n
12
29
647
716
583
587
%
VUB03KD SC2
VUB03TG SC2A
VUB03TG SC2B
579
621
631
VUB03TG
SC2A
SC2B
21
25

**Figure S11. Methylation pattern up- and downstream of the CTG repeat from *MSH2* transgene VUB03-DM1 bulk cell lines.** *MSH2* was re-introduced in VUB03KD-SC1, VUB03KD-SC2 and VUB03KD-SC3 and a bulk sample was collected before clonal expansion to asses methylation status before cloning. The upstream methylation is shown for 25 CpG sites and for the downstream site, 11 CpG sites were analyzed. A total of 100 epi-alleles were analyzed per bulk sample. The bubble size indicates the percentage of epi-alleles with that specific amount of CpG methylation. Abbreviations: KD: *MSH2* knock-down, TG: *MSH2* transgene expression

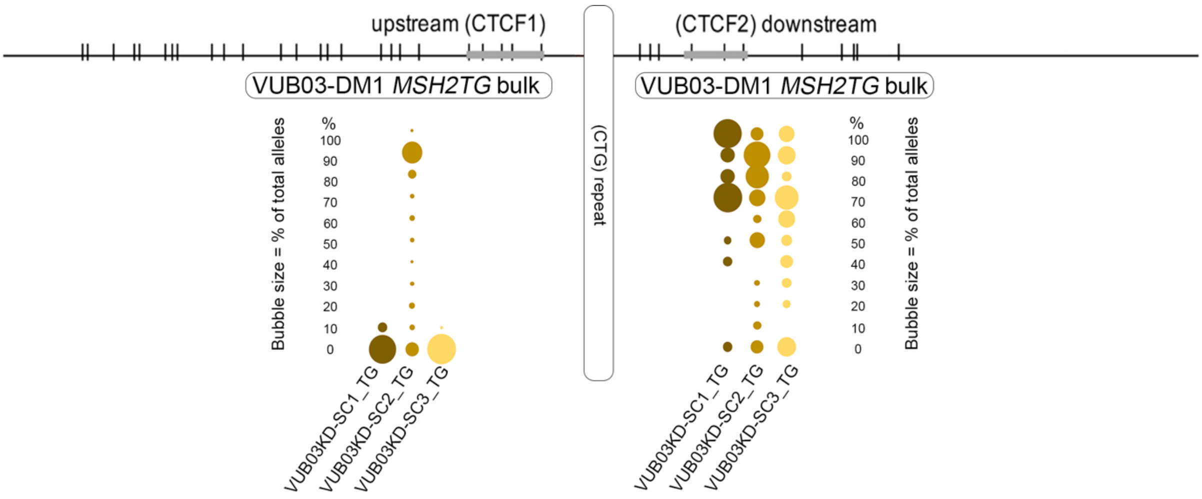

**Table S7. Kolmogorov Smirnov test output on the CpG methylation data upstream of the repeat in MSH2KD compared to MSH2KD_TG samples.** Samples were compared at passages 12 before transgene expression and passage 19 after transgene expression but before clonal expansion.

| VUB03KD-SC1 - VUB03KD-SC1_TG | VUB03KD-SC2 - VUB03KD-SC2_TG | VUB03KD-SC3 - VUB03KD-SC3_TG |
|---|---|---|
| **p12 - p19 bulk** | **p12 - p19 bulk** | **p12 - p19 bulk** |
| **p=0,076** | **p=0,461** | **p=0,076** |
| data distribution is the same | data distribution is the same | data distribution is the same |
